# Frontal Eye Field Leads a Distributed Oculomotor Circuit for Abstract Categorical Decisions

**DOI:** 10.64898/2026.06.30.735630

**Authors:** Ou Zhu, Vinay Shirhatti, Maura Garza, Yunlong Xu, Samuel David, Chris K. Hauser, Brent Doiron, David J. Freedman

**Affiliations:** Department of Neurobiology, The University of Chicago, Chicago, IL 60637, USA; Neuroscience Institute, The University of Chicago, Chicago, IL 60637, USA

## Abstract

Flexible decisions require the brain to transform sensory evidence into abstract, task-relevant variables and then into actions. Understanding this process requires identifying how distributed neural populations represent sensory, cognitive, and motor variables, and how interareal interactions mediate transformations between them. We simultaneously recorded population activity in frontal eye field (FEF), lateral intraparietal area (LIP) and superior colliculus (SC) while monkeys performed a flexible yet urgent visual motion-categorization task. Within this network, FEF first encoded abstract categories, followed by SC and then LIP. LIP showed the earliest encoding of visual stimulus features, but a later encoding of upcoming saccades. Single-trial analyses revealed directed information flow from FEF to LIP populations for category-and choice-related signals. Reversible FEF inactivation impaired categorization and saccadic choice, causally implicating FEF in category-guided action. These findings reveal a differentiated FEF-LIP-SC circuit for transforming sensory evidence into abstract categorical decisions and the actions used to report them.

## Introduction

Humans and non-human primates have a remarkable capacity to make flexible, yet rapid, context-dependent decisions about incoming visual stimuli. Visually-based abstract decisions engage a broad hierarchy of cortical and subcortical brain regions which analyze visual stimulus features, compute the significance or meaning of those stimuli, and select context-appropriate motor responses^1,2^. There is widespread agreement that primary sensory and primary motor regions are critical sites of sensory and motor processing, respectively. Less clear is how distributed circuits implement flexible decisions: how sensory representations are transformed into abstract, task-relevant variables, how these variables are coordinated across brain regions, and how they are converted into context-appropriate actions. A traditional modular view of functional localization would ascribe cognitive functions to associative/executive regions, such as the prefrontal cortex (PFC) and posterior parietal cortex (PPC). However, the advent of increasingly sophisticated neural recording approaches for comparing population activity across multiple brain regions has revealed that task-related functions can be encoded and even causally mediated by a diverse set of brain regions extending into areas traditionally considered to be primarily sensory or motor^3^ (also see ^4,5^).

A key challenge is to determine how these distributed representations are communicated and coordinated across interconnected neural populations. Abstract categorical decisions provide a powerful framework because they require sensory evidence to be transformed into a learned, task-relevant variable and then into an action. Previous work has shown that category information is represented across multiple cortical and subcortical regions, including prefrontal, parietal, and oculomotor structures. These findings argue against a strictly modular view in which abstract decisions are computed only in canonical association cortex and merely read out by downstream motor systems. However, they leave unresolved how sensory, categorical, and action-related signals are coordinated within a brain-wide circuit during decision formation.

The primate visual system and oculomotor network provide a well-defined and tractable circuit for addressing this question. Prior work using learned visual categories has traced visual and category-related signals across a network primate visual, associative, and oculomotor areas, including the middle temporal (MT) and medial superior temporal (MST) areas, LIP, medial intraparietal (MIP) area, lateral prefrontal cortex (LPFC), and SC^6,7,8,9,10,11,12,13,14^. FEF, LIP, and SC are strongly interconnected and have well-established roles in visual selection, spatial attention, and gaze control^15,16,17^. Furthermore, LIP and SC have been shown to encode learned visual categories and to contribute causally to abstract categorization^18,19,20^, including in tasks that dissociate categorization from their classic roles in saccade generation. These observations raise a broader possibility: rather than serving primarily as a sensorimotor readout pathway, the FEF-LIP-SC network may participate directly in transforming sensory evidence into abstract decisions and the actions used to report them. Testing this possibility requires simultaneous population recordings across the circuit, because recordings from each area separately cannot determine the temporal ordering or directed interactions among areas on individual trials.

We therefore recorded simultaneously from neuronal populations in FEF, LIP, and SC while macaque monkeys performed a visual-motion-based rapid categorization (RC) task adapted from compelled-saccade decision paradigms^21,22^. The task dissociated abstract category from the saccade used to report it by randomizing the locations of red and green choice targets across trials, and used complementary spatial configurations of motion stimuli and saccade targets to separately emphasize and examine sensory-processing and motor-preparation components of the decision process. This design allowed us to measure how behavior transitioned from guessing to informed categorical choices as a function of available stimulus evidence, while also dissociating motion direction, abstract category, target configuration, and the saccade used to report the decision. This design enabled us to compare when sensory, categorical, and motor variables emerged across the three areas, to assess directed interareal interactions at the single-trial level, and to test the causal role of FEF using reversible inactivation.

Here we show that FEF, LIP, and SC instantiate a functional cortical–subcortical circuit for transforming sensory evidence into abstract categorical decisions and the saccades used to report them. Neural activity in all three areas encoded visual motion direction, abstract stimulus category, target configuration, and saccade direction, but these signals emerged with distinct temporal profiles. LIP provided the earliest representation of visual stimulus features and target configuration, whereas FEF showed the earliest encoding of abstract category and, together with SC, the earliest encoding of impending saccadic choice. Single-trial analyses of simultaneously recorded populations revealed directed information flow from FEF to LIP for category-and choice-related signals, and from FEF to SC for category-related signals. Finally, reversible inactivation of FEF impaired rapid categorization and saccadic choice, demonstrating a causal contribution of FEF to the decision-to-action transformation. Together, these findings argue that the FEF-LIP-SC network is not merely a downstream readout pathway, but participates directly in transforming sensory evidence into abstract categorical decisions and actions.

## Results

### Task design and behavioral performance

We trained two male rhesus monkeys (M and S) to perform a RC task that adapts a compelled-saccade paradigm to study abstract visual motion categorization under time pressure^21^. On each trial, the animals fixated a central point while first a random-dot motion stimulus and then two colored saccade targets (red and green) appeared in the periphery (Fig. 1A, Fig. S1A). This initial random-dot stimulus was 100% noise (i.e. had no net motion direction), and therefore uninformative about the motion category. After a random interval of 250-450 ms the fixation point disappeared, serving as a go cue for the monkeys to report their decision about motion category with a saccade to the corresponding color target within 500 ms. Importantly, the coherent motion stimulus, required to make an informed decision, was made available to the monkey on each trial at a variable time with respect to the go cue. Following a random interval (in the range of 0 to 250 ms w.r.t the go cue in Monkey M, and similarly-100 to 100 ms in Monkey S; see Methods for details) the noise stimulus was replaced by a 100% coherent motion stimulus moving in one of six evenly spaced directions. The six motion directions were arbitrarily divided into two 180°-wide experimenter-defined categories established through training, with three directions in each (Fig. 1A inset; Fig. S1A inset). The color identity (red vs. green) of the two targets were randomized from trial to trial (i.e. left-right randomized in the horizontal, and up-down randomized in the vertical task orientation), allowing us to dissociate the category of the stimulus from the direction of the saccade used to report that category.

**Figure 1:**
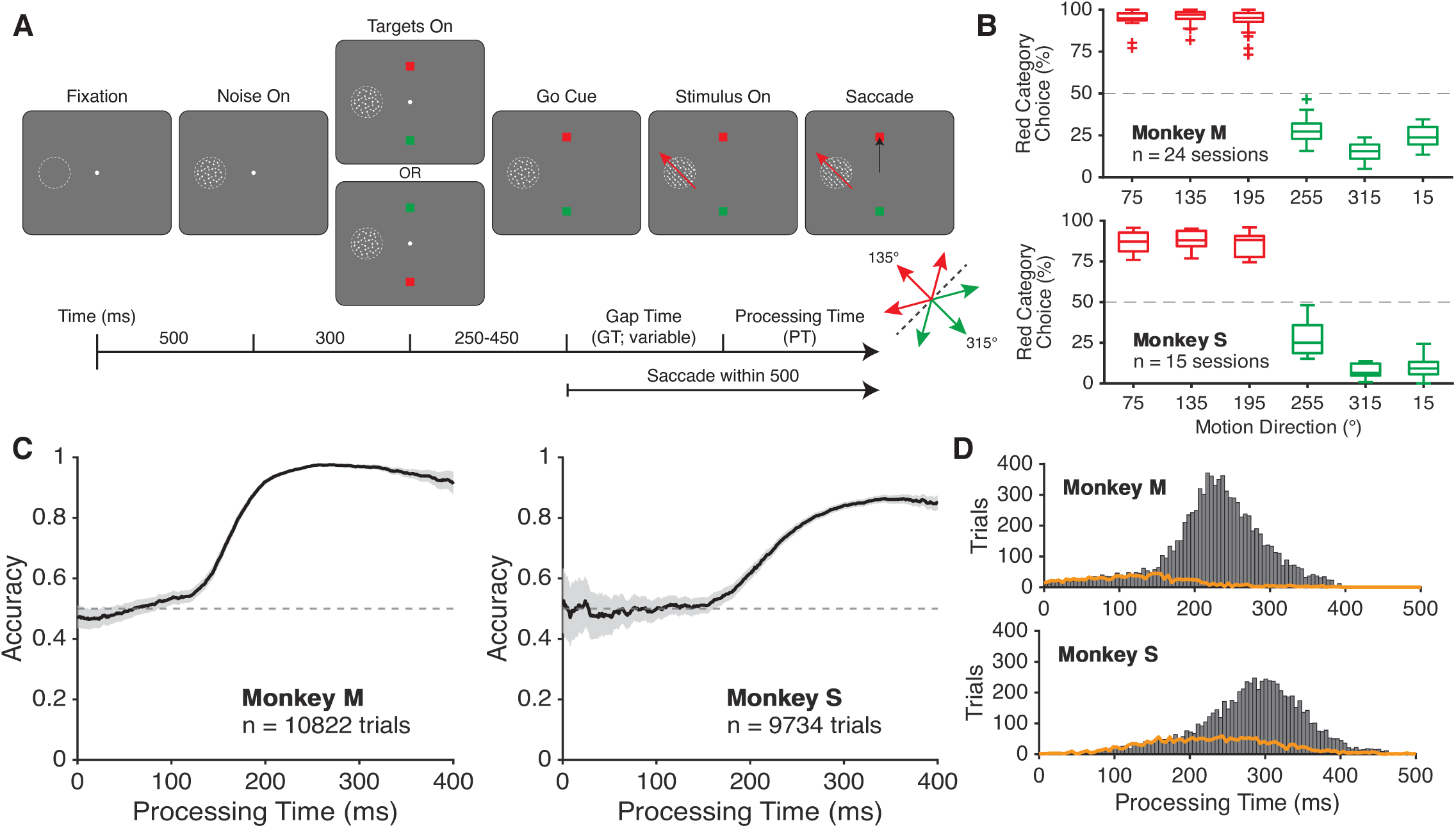
Task outline and behavioral analyses for monkeys performing the rapid categorization task with vertical targets. **(A)** Trial structure of the rapid categorization task, such that the motion stimulus is positioned in the receptive field of recorded neurons (left visual field). Monkeys initiate each trial with central fixation and maintain fixation as motion stimuli and saccade targets appear in the periphery. After the fixation point disappears (Go Cue), the monkey must saccade to a color target within 500ms. However, the motion stimulus only becomes informative after a variable Gap Time (GT) following the Go Cue (see Methods). The positions of the color targets are randomized every trial. The category identity of the six motion directions are shown in the bottom-right inset, with center directions labeled. **(B)** Monkeys’ behavioral performance for each direction across all recording sessions, across all processing times. **(C)** Monkeys’ categorization accuracy quantified across a range of processing times (PT); accuracy quantified as the fraction of trials the monkey responded correctly. Shaded areas indicate SEM calculated from binomial statistics (see Methods). **(D)** Distribution of processing times of correct (gray bars) and incorrect trials (orange lines) from both monkeys.

Enforcing a variable gap time between go cue and coherent motion stimulus onset produces a continuum of processing times (PT), defined as the interval between the onset of the 100% coherent motion and the onset of the animal’s saccade, during which sensory evidence is available for decision-making (Fig. 1A, Fig. S1A). The task design incentivizes the monkeys to report their decisions rapidly. Short PTs require the animal to respond under severe uncertainty, with little to no information about motion direction, whereas longer PTs allow for reliable perception of motion direction. As in previous compelled-saccade studies, this design yields a systematic transition from guessing to informed choices as a function of PT and thus offers a behavioral benchmark for interpreting the timing of neural encoding across areas.

Neuronal recordings were performed once monkeys were fully trained on the task, with high overall accuracy at long processing times. Psychometric functions measured across recording sessions revealed clear categorical behavior: performance was similar for motion directions within the same category and changed sharply at the category boundary, for both vertical-and horizontal-target orientations (Fig. 1B; Fig. S1B). Crucially, accuracy increased monotonically with PT: at very short PTs, performance was near chance, reflecting decisions based on little (or even zero) evidence, whereas at longer PTs (∼200–300 ms) both animals approached asymptotic accuracy on the categorization task (Fig. 1C; Fig. S1C). Errors were disproportionately concentrated at short PTs, as reflected in the PT distributions of correct and incorrect trials (Fig. 1D; Fig. S1D).

These behavioral patterns demonstrate that the monkeys solve the RC task by making rapid, abstract categorical decisions with accuracy predictably depending on the time available to accumulate motion evidence. The PT-dependent transition from chance to reliable categorization performance provides a natural timescale for interpreting when LIP, FEF, and SC begin to encode motion direction, stimulus category, target configuration, and saccade direction in the neural analyses that follow.

### Simultaneous population recordings in FEF, LIP, and SC

To compare stimulus-, decision-, and saccade-related activity across the oculomotor network, we recorded simultaneously from neural populations in FEF, LIP, and SC in the two male rhesus monkeys trained to perform the RC task (M and S). Each animal was implanted with two recording chambers in the right hemisphere: one giving access to FEF and a second allowing targeting of both LIP and SC, guided by pre-implant structural MRI and CT imaging. Recording sites in all three areas were selected based on the presence of visual receptive fields and/or saccade-related response fields (RFs) in the left visual hemifield, identified with a memory-guided saccade (MGS) task in each session.

During performance of the RC task, we used up to six linear multichannel arrays per session (dense 24-or 32-channel probes; typically two per area; see Methods) to record spiking activity within each region (Fig. 2A, 2B). Across all recording sessions, we obtained well-isolated single-unit activity from 1,209 FEF neurons (Monkey M: 595; Monkey S: 614), 1,165 LIP neurons (Monkey M: 672; Monkey S: 493), and 520 SC neurons (Monkey M: 253; Monkey S: 267). We also obtained a population of multiunit clusters (defined by Kilosort spike sorting, see Methods) from FEF (Monkey M: 417; Monkey S: 292), LIP (Monkey M: 631; Monkey S: 214), and SC (Monkey M: 240, Monkey S: N=118). In the current study, measures of single neuron activity and selectivity (e.g. category tuning index) were applied across single units only, whereas population measures (e.g. category and saccade decoders) were run across combined populations of single and multiunits. However, qualitatively similar results were found when analyses were restricted to only single units. These large, simultaneously recorded populations allowed us to quantify not only the strength and timing of task-related visual, cognitive, and saccadic encoding within each area, but also the temporal ordering and putative information flow across areas within individual sessions.

**Figure 2:**
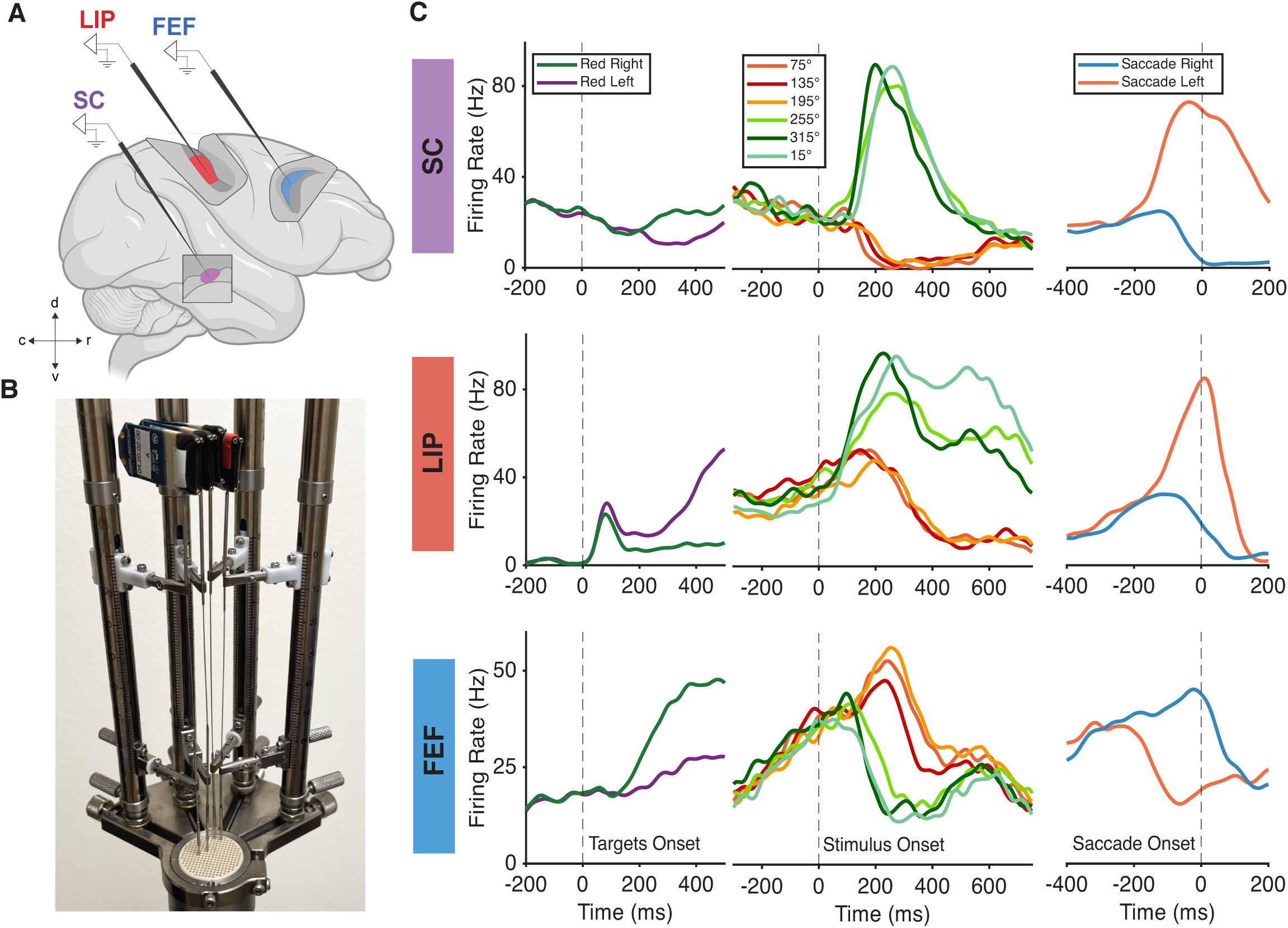
Single neurons in LIP, FEF, and SC are modulated by target identities, motion categories, and saccade directions. **(A)** Schematic of recorded brain locations. (B), Picture demonstrating four linear arrays (Plexon S-probes) simultaneously positioned within one recording grid to target areas LIP and SC with two arrays each. (C), Example neurons from each area that were simultaneously selective for visual motion category, visual motion direction, saccade direction, and target configuration, with firing rates aligned to three separate event times: the onset of the targets, the onset of the full coherence motion stimulus, and the onset of the animal’s saccade. Each example neuron shown was determined to be selective for task features and saccade direction by an ANOVA model fit at four time points relative to stimulus onset in at least one of the task orientations (p<0.05/8). Firing rates were trial-averaged by the feature most relevant for each alignment time, e.g. target color in receptive field (left panels), motion direction (middle panels), and saccade direction (right panels). Horizontal-orientation trials were used for trial-averaging by target configuration or saccade direction (left and right panels, respectively), and vertical-orientation trials filtered to have ≥200ms processing time for motion direction (middle).

We first asked, at a coarse level, to what extent single neurons in each area were modulated by the main task variables. Using a nested ANOVA on firing rates aligned to stimulus onset, we assessed selectivity for motion category, motion direction, saccade direction, and target configuration (the red/green target positions–e.g. in horizontal-target orientation trials, whether the green target was on the right or left). Over 80% of neurons in each region were significantly modulated by at least one of these features (Table S1A), with more than two-thirds significantly modulated by two or more features (Fig. S2). Up to 45% of neurons in FEF, LIP, and SC showed robust modulation by all four task variables (Fig. S2). To illustrate this mixed selectivity, Fig. 2C shows example neurons from LIP, FEF, and SC recorded during the RC task. Each example neuron exhibits clear modulation by target configuration (left panels), motion category and/or motion direction (middle panels), and saccade direction (right panels), demonstrating that individual neurons in all three areas can concurrently encode stimulus, context, and choice variables.

This combination of simultaneous multi-area recordings and strong multi-feature modulation at the single-neuron level provides the foundation for the analyses that follow. In the next sections, we use population-level decoding and latency analyses to dissect how neuronal encoding of abstract category, motion direction, saccade direction, and target configuration emerge over time across FEF, LIP, and SC, and how these dynamics relate to the animals’ rapid categorical choices.

### Stimulus-related encoding: FEF leads category encoding, LIP/SC leads direction encoding

We next quantified how information about the motion stimulus is distributed across LIP, FEF, and SC at the population level. We focused on two key aspects of stimulus encoding: the raw motion direction of the stimulus and its abstract category. To isolate stimulus-related responses, these analyses used correct trials from the stimulus and target configuration in which the motion stimulus was presented in the neurons’ contralateral visual field and the saccade targets were positioned vertically. The distribution of neurons’ RFs is shown in Fig. S3. We also focused on trials with long processing times (PT ≥ 200 ms) to ensure that sufficient motion information was available before the saccadic choice.

To characterize category encoding in each area, we first computed a receiver-operating characteristic (ROC)-based category tuning index (rCTI)^23^ over time for neurons that were significantly direction-or category-selective as determined by the nested ANOVA. Population-averaged rCTI increased after motion onset in all three regions (Fig. 3A; see Methods), which is expected given that many individual neurons show robust binary-like selectivity for one motion category over the other (Fig. 2C). We then trained linear support vector machine (SVM) decoders on pseudopopulations of single and multiunits constructed separately for each brain area and monkey to predict the stimulus category from time-resolved spike counts, using a decoding approach which assesses category encoding independent of direction tuning (Fig. 3B; Table S2A; see Methods). Category decoding accuracy rose rapidly after stimulus onset in all three areas, but with a consistent temporal ordering in both monkeys: decoding emerged earliest in FEF, followed by SC then LIP. FEF decoders achieved high accuracy within ∼100–150 ms of stimulus onset, whereas LIP and SC decoders crossed the same accuracy criterion significantly later. The differences in selectivity latency (see Methods) supporting FEF as the site of earliest category encoding were found to be significant by permutation tests comparing the distribution of decoder latencies across areas (Fig. 3B, right; Table S2A). Notably, this order of encoding was consistent across target orientations; FEF leads SC, then LIP, regardless of whether the motion stimulus or a saccade-target was placed in the left visual hemifield (Fig. 3B,S4A; Table S2A,S3A).

**Figure 3:**
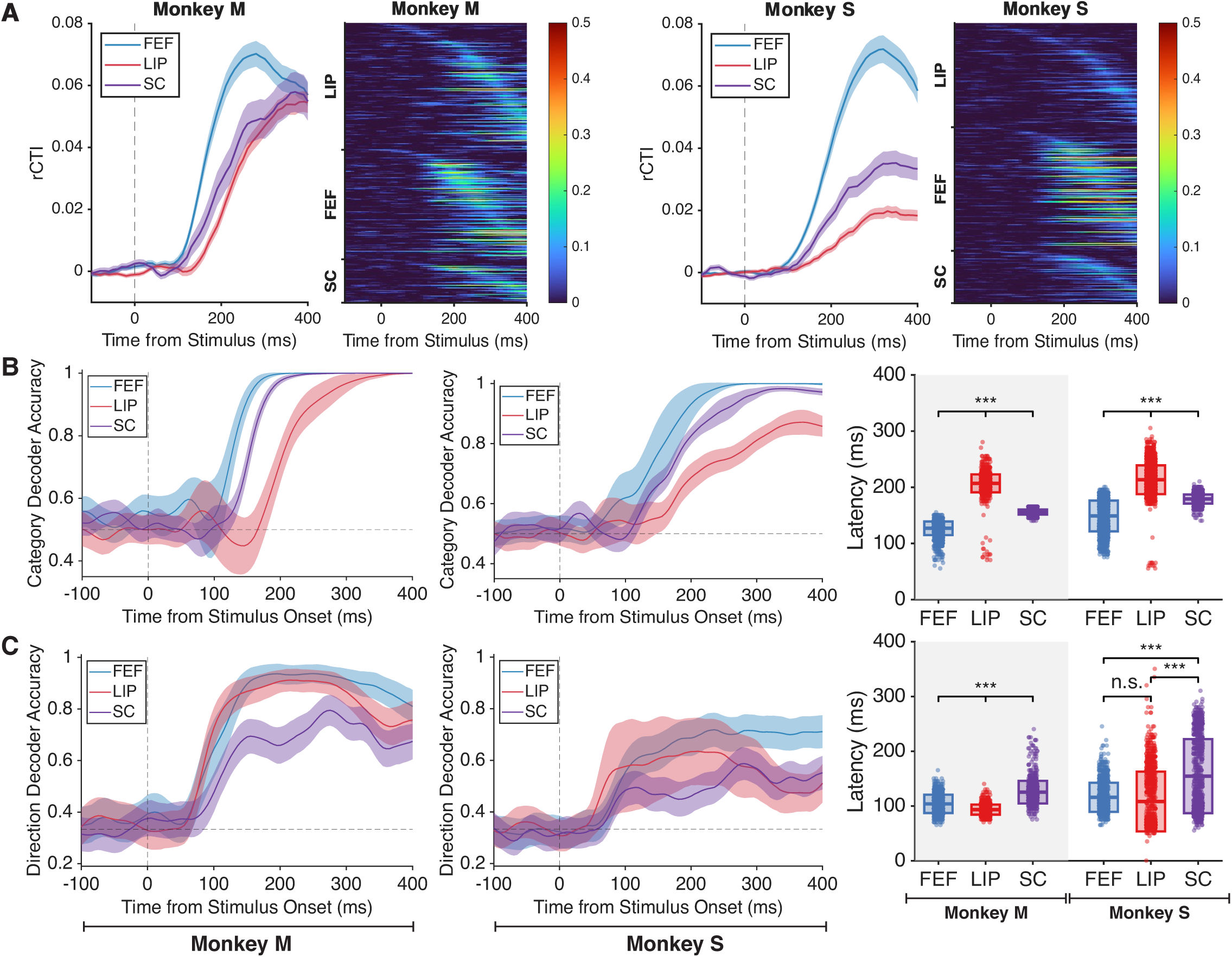
Direction and category selectivity are distributed at different time scales across LIP, FEF, and SC. (A), Average single neuron ROC-based category tuning index (rCTI) by brain region (far left panel) and individual single neuron rCTIs arranged in rows as a heatmap (middle left panel) for Monkey M. The same format is used on the right panels to display results of single neurons for Monkey S. Only stimulus (category and direction) selective neurons as determined by ANOVA are shown. Shaded areas indicate SEM across single neurons. **(B),** Accuracy of linear support vector machines (SVMs) trained to predict the category of stimuli using activity from pseudopopulations constructed for each area. Left panel denotes the time course of SVM accuracy for Monkey M. Middle panel denotes the time course for Monkey S. Correct vertical-target orientation trials in which the animal had at least 200ms of processing time, and the motion stimulus was placed to the left of fixation were used for these analyses. Shaded intervals represent ± one standard deviation across pseudopopulations. Right panel shows scatter plots of the latency of SVMs for each pseudopopulation, defined as the first time point the SVM crossed an accuracy threshold of halfway between chance and the maximum accuracy attained by the SVM, for five consecutive time points (25ms). Box plots denote mean ± one standard deviation. Gray shading denotes neurons from Monkey M. **(C),** The same layout as **b,** but displaying results for SVMs trained to predict the direction of motion stimuli using activity of pseudopopulations for each area. *** denotes p<0.001, subsampled permutation test.

We applied a similar pseudopopulation decoding approach to assess motion-direction encoding independent of category encoding (see Methods), training SVMs to decode the direction of motion (within a category) from population activity (Fig. 3C; Table S2B). In contrast to the category results, direction decoding showed a different temporal hierarchy: motion-direction information emerged earliest in LIP, followed by FEF, and latest in SC. Note that in Monkey M, LIP significantly leads both FEF and SC. For Monkey S, while the temporal ordering remains consistent with Monkey M, the latency differences between FEF and SC did not reach significance. This ordering is consistent with LIP’s known role as a direct recipient of motion signals from dorsal-stream visual areas (e.g. MT and MST), and suggests that within this network LIP provides the earliest access to raw visual motion information. Surprisingly, in the horizontal-target task orientation, in which a saccade-target was placed in the left visual hemifield, SC had the earliest motion direction encoding latency (Fig. S4B; Table S3B), suggesting an additional role for SC in the motion processing of the RC task. It is possible that the difference in SC and LIP motion direction encoding latencies across the task orientations is due to differences in receptive field concentrations among the populations recorded (Fig. S3); however, such differences did not impact categorization latencies.

For the decoding analyses of category and direction, we examined the robustness of our latency measurements as a function of neural population size in each of the brain areas. We did so in order to confirm that our latency results–specifically the ordering of selectivity latencies across FEF, LIP, and SC–were not critically dependent on the number of neurons we happened to have collected in our experiments. To do so, we repeated the decoding and latency analyses by subsampling units across a range of population sizes ranging from 50 to the minimum available number of units across the three areas. These results, shown in Fig. S5A,B, strengthen and confirm results in Fig. 3B,C. Across all pseudopopulation sizes tested in both monkeys in the vertical-target task orientation, mean category encoding latency followed the order of FEF-SC-LIP, with latency differences becoming more apparent at higher pseudopopulation sizes in Monkey S (Fig. S5A). Similarly, the LIP-FEF-SC order of mean latencies in the vertical-target task orientation was consistent across all pseudopopulation sizes tested in Monkey M (Fig. S5B, left). In Monkey S, LIP and FEF mean latencies consistently led SC across pseudopopulation sizes, although LIP did not consistently lead FEF (reflective of nonsignificant differences in Fig. 3C; Fig. S5B, right). Results were likewise robust across population sizes in the alternative, horizontal-target task orientation (data not shown).

Together, these analyses reveal a dissociation between the latencies of category and direction encoding across the three regions. LIP/SC leads the encoding of raw motion direction, trailed by FEF, but FEF leads the encoding of abstract motion category, trailed by SC and LIP. This pattern is difficult to reconcile with a model in which category representations are computed solely in LIP or SC and simply inherited by FEF. Instead, it suggests that FEF may be a key locus for transforming upstream motion information into abstract categorical representations that are then broadcast to SC and fed back to LIP. In the next section, we turn to saccadic-choice and motor-related encoding, examining how neural encoding of target configuration and saccade direction emerge across the same three areas.

### Target selection and saccade encoding: LIP and SC lead in target configuration, FEF and SC lead in saccade direction

We next examined how FEF, LIP, and SC encode variables related to target selection and the planning and execution of saccades. We focused on two main factors: the target configuration (the spatial arrangement of the red and green saccade targets) and the direction of the saccade used to report the decision. For these analyses, we focused on trials in which the saccade targets were positioned along the horizontal axis, so that the targets were in the contralateral and ipsilateral visual hemifields, respectively. As for the stimulus-related analyses, we used SVM-based pseudopopulation decoding applied separately to each brain area and monkey, allowing us to assess the strength and timing with which each area encodes the identity (red vs. green) of the target in the contralateral hemifield or the direction of the saccade (contraversive vs. ipsiversive). For target configuration decoding, neural activity was aligned to the onset of the red and green targets, and for saccade direction decoding, to the time of saccade onset. Only correct trials were used to decode the configuration of saccade targets, whereas both correct and incorrect completed trials were included for decoding the direction of the impending saccade..

Target configuration information was decodable in all three regions, but its temporal evolution differed systematically across areas. In both animals, target decoder accuracy rose above chance earliest in LIP, followed by SC, and latest in FEF (Fig. 4A, left and middle; Table S4A), with statistically significant latency differences between regions in both monkeys (subsampled permutation tests on pseudopopulation latency distributions; Fig. 4A, right; Table S4A). Thus, at the population level, LIP provides the earliest representation of which color target is located in the contralateral visual field, consistent with a role in encoding visual features of task-relevant saccade targets. In the vertical-target orientation, in which the motion stimulus, instead of a saccade target, is positioned in the contralateral visual field, LIP and SC consistently encode the (up/down) spatial configuration of targets before FEF, as in the horizontal orientation, although SC slightly but significantly leads LIP (Fig. S6A; Table S5A).

**Figure 4:**
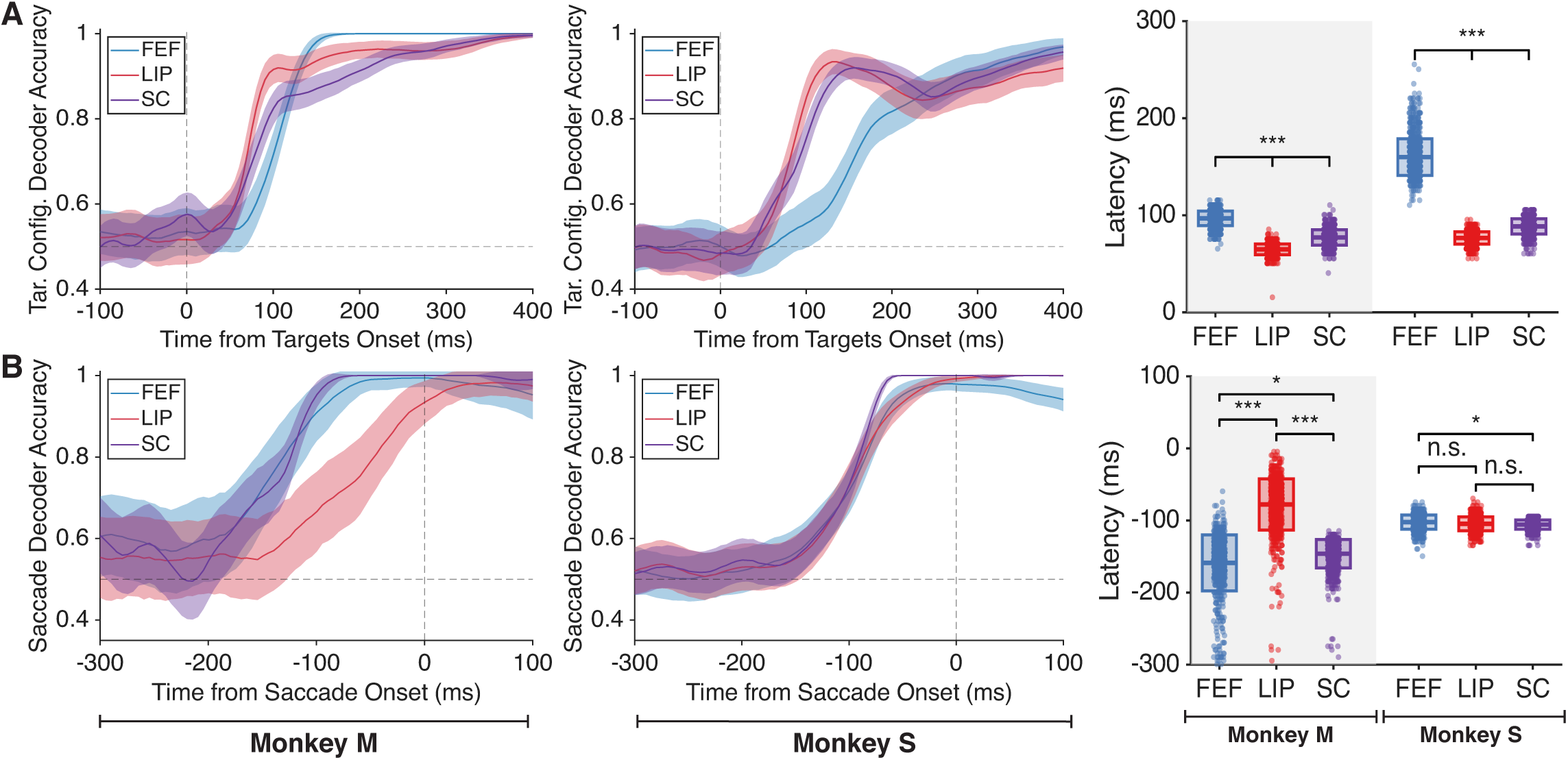
LIP leads the encoding of target configuration while core oculomotor output areas lead the encoding of saccade direction. (A), Accuracy of linear support vector machines (SVMs) trained to predict the target configuration (e.g. target color in RF) using activity from pseudopopulations constructed for each area. Left panel denotes the time course of SVM accuracy over time for Monkey M. Middle panel denotes the time course for Monkey S. Correct trials of the horizontal orientation in which one target was placed to the left of fixation were used. Shaded intervals represent ± one standard deviation across pseudopopulations. Right panel shows scatter plots of the latency of SVMs for each pseudopopulation, defined as the first time point the SVM crossed an accuracy threshold of halfway between chance and the maximum accuracy attained by the SVM, for five consecutive time points (25ms). Box plots denote mean ± one standard deviation. Gray shading denotes neurons from Monkey M. **(B),** The same layout as **a**, but displaying results for SVMs trained to predict the monkey’s saccade direction from both correct and incorrect trials, using activity of pseudopopulations for each area. *** denotes p<0.001, ** denotes p<0.01, * denotes p<0.05, subsampled permutation test.

All three regions encoded the direction of the impending saccade prior to its execution. In both monkeys and versions of the task (vertical-and horizontal-target orientations), encoding of saccade direction primarily emerged earlier in FEF and SC compared to LIP (Fig. 4B, Fig. S6B; Table S4B, S5B). The only exception to this finding was the result in Monkey S during the horizontal-target orientation of the task, in which the latency of encoding the upcoming saccade in LIP was not significantly different from the latencies in FEF and SC (Fig. 4B, center and right panels; Table S4B).

As we did for the decoding latencies for category and motion direction, we examined the robustness of these latency measurements as a function of neural population size in each of the brain areas (Fig. S5C,D). For target configuration encoding in the horizontal-target task orientation, the mean encoding latency in LIP consistently leads SC and FEF in both monkeys across all pseudopopulation sizes tested (Fig. S5C). Further, the full LIP-SC-FEF order is consistent across all pseudopopulation sizes in Monkey S. For saccade encoding in the horizontal-target task orientation, FEF and SC consistently lead LIP across all pseudopopulation sizes tested in Monkey M (Fig. S5D, left; agreeing with Fig. 4B, left). For the alternative, vertical-target task orientation, average target configuration encoding latencies in LIP and SC consistently lead FEF across all pseudopopulation sizes tested in both monkeys, while FEF and SC consistently lead LIP in mean saccade direction encoding latencies (data not shown).

These findings reveal a functional dissociation within the oculomotor network. LIP and SC provide the earliest representation of which color target occupies the contralateral visual field, consistent with a prominent role in encoding task-relevant visual and spatial context. The core oculomotor output areas (FEF and SC) primarily provide the earliest representation of the saccadic choice, paralleling their lead in abstract category encoding and supporting a central role in initiating the motor plan for reporting the animal’s categorical decision.

### Single-trial analyses reveal interareal dynamics within the FEF-LIP-SC circuit

Our decoding analyses that were based on constructing pseudopopulations indicate shorter-latency emergence of abstract category signals in FEF relative to SC and LIP, and saccadic-choice signals in FEF and SC relative to LIP. However, pseudopopulation analysis does not preserve trial-to-trial neuronal covariation within or across simultaneously recorded areas. Thus, any conclusions based on pseudopopulations ignore within trial variability, yet animal decisions are necessarily based on within trial neuronal responses. Further, while the neural computations are distributed across multiple brain areas, pseudopopulation analysis is blind to potential between area coordination that would point to how category or saccade information flows within the full circuit. We therefore took advantage of our simultaneous population recordings across all three areas to test for directed coupling between FEF, LIP, and SC at the single-trial level (Fig. 5).^24,25^

**Figure 5:**
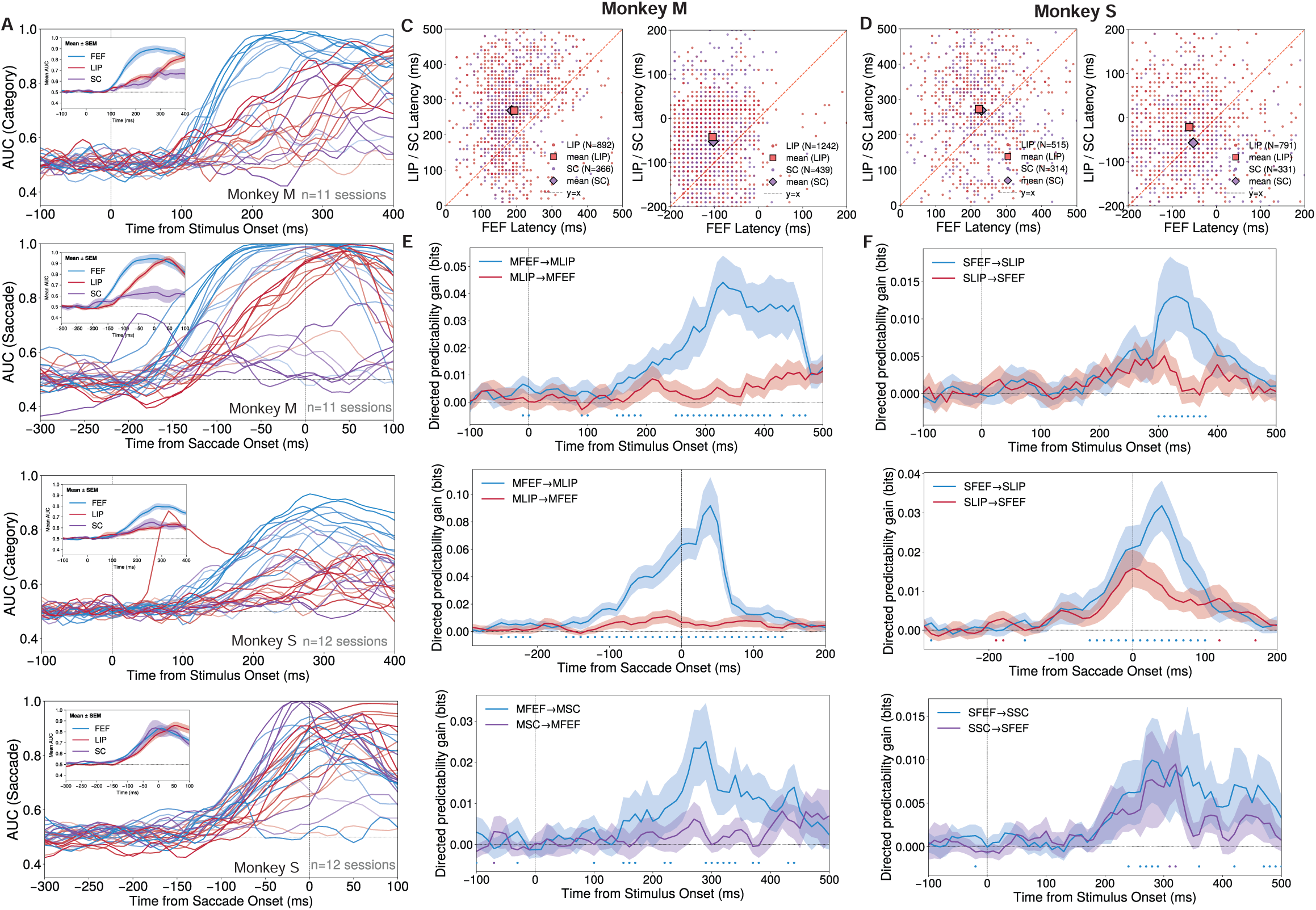
Single-trial analyses reveal interareal dynamics within the FEF-LIP-SC circuit. Category analyses are stimulus-aligned (vertical-target trials). Saccade analyses are saccade-aligned (horizontal-target trials). **(A,C,E)** Monkey M. **(B,D,F)** Monkey S. **(A,B)** Time-resolved decoding performance (AUC) measuring how well activity projected onto session-specific encoding axes discriminates between conditions (top: category; bottom: saccade direction). Each thin trace is one session. Within each color, trace opacity is scaled by the session’s peak AUC: sessions with higher peak decoding performance are rendered more opaque, and those with lower peak performance more transparent. Insets show the across-session mean ± SEM. Colors: FEF (blue), LIP (red), SC (purple). Dashed line: alignment event (stimulus onset for category; saccade onset for saccade direction). **(C,D)** Single-trial onset latencies comparing FEF versus LIP (red points) and FEF versus SC (purple points) for category (left) and saccade direction (right). Onset is defined as the first time signed evidence exceeded a baseline-derived threshold (μ + 4σ) for 5 consecutive bins; only trials with defined onsets in both areas of a pair were included. Dashed line: y = x. Large markers indicate the mean latency for each area pair. Mean latency differences (partner − FEF) were: Monkey M category: LIP−FEF = 75.6 ms (N = 892, p < 0.001), SC−FEF = 83.2 ms (N = 366, p < 0.001); saccade: LIP−FEF = 61.8 ms (N = 1,242, p < 0.001), SC−FEF = 50.1 ms (N = 439, p < 0.001). Monkey S category: LIP−FEF = 50.1 ms (N = 515, p < 0.001), SC−FEF = 39.5 ms (N = 314, p < 0.001); saccade: LIP−FEF = 40.2 ms (N = 791, p < 0.001), SC−FEF =-4.4 ms (N = 331, p =0.489). P-values were assessed with sign-flip permutation tests on paired latency differences. **(E,F)** Directed predictability gain (bits): improvement in predicting one area’s current feature evidence from the other area’s recent history (80 ms lag window for FEF–LIP category, 50 ms for FEF–LIP saccade, 30 ms for FEF–SC) beyond what can be predicted from the target area’s own history. Top: category, FEF–LIP. Middle: saccade, FEF–LIP. Bottom: category, FEF–SC. Curves show mean ± SEM across sessions; line color indicates the source area (blue: FEF→target, red: LIP→target, purple: SC→target). Colored dots mark time bins where one direction significantly exceeded the reverse under a stratified trial-shuffle null (p < 0.05); dot color indicates which direction was larger. The null distribution was generated by shuffling trial identities within task-condition strata, thereby preserving condition-specific evoked structure while breaking trial-by-trial pairing between areas.

We first focused on FEF and LIP, which showed reliable differences in average encoding latency at the pseudopopulation level for both category and saccade direction. For each session and area, we identified a linear decoding axis that best discriminates trials by the relevant task variable (e.g. category or saccade direction, see Methods). We then projected each trial’s population activity onto this axis to obtain a one-dimensional “feature evidence” trace that tracks the strength of encoding over time within each area. To assess how well these axes captured the relevant signals, we computed time-resolved decoding performance (AUC) by measuring how well the projected activity discriminated between the two categories (or saccade directions) at each time point (Fig. 5A,B; insets show the across-session mean). Each trace represents one recording session; sessions were included in subsequent analyses if peak AUC reached 0.65 in each of the areas since we reasoned that without reliable encoding in both areas, the analysis of cross-area covariation is not well defined. For category encoding (Fig. 5A,B, top), this yielded 11 sessions for Monkey M and 12 for Monkey S; for saccade direction encoding (Fig. 5A,B, bottom), this yielded 11 sessions for Monkey M and 11 for Monkey S. Satisfyingly, in both monkeys and for both variables, decoding performance rose earlier in FEF than LIP for most sessions–mirroring the results from the pseudopopulation analysis (Figs. 3-4).

We next compared single-trial onset latencies between areas by identifying when each trial’s feature evidence first crossed a trial-specific detection threshold (Fig. 5C,D). For each trial, we defined onset as the first time the signed evidence exceeded a baseline-derived threshold (μ + 4σ) for five consecutive bins. FEF systematically preceded LIP for both task variables in both animals. For category (Fig. 5C,D, left; red points), FEF led LIP by a median of 80.0 ms in Monkey M (mean: 75.6 ms; N = 892 trials; p < 0.001, sign-flip permutation test) and 60.0 ms in Monkey S (mean: 50.1 ms; N = 515 trials; p < 0.001). For saccade direction (Fig. 5C,D, right; red points), FEF led by a median of 70.0 ms in Monkey M (mean: 61.8 ms; N = 1,242 trials; p < 0.001) and 40.0 ms in Monkey S (mean: 40.2 ms; N = 791 trials; p < 0.001), consistent with the pseudopopulation results (Figs. 3–4).

It is unclear whether trial-to-trial fluctuations in feature evidence are coupled between FEF and LIP (Fig. 5E,F, top and middle). To explore this possibility we used lagged ridge regression to measure how well one area’s recent history predicts the other area’s current state, beyond what can be predicted from the target area’s own history. We call this improvement “directed predictability gain”, which quantifies how much predictive information the source area provides about the target area. We computed this for both directions (FEF→LIP and LIP→FEF) and assessed significance using a permutation test that shuffled trial identities within condition strata, thereby preserving condition-specific evoked structure while breaking trial-by-trial pairing between areas. FEF→LIP gain was consistently larger than LIP→FEF, with significant positive net flow (defined as the FEF→LIP gain minus the LIP→FEF gain; p < 0.05, blue dots) during 140–190 ms and 250–400 ms post-stimulus for category in Monkey M (Fig. 5E, top), 300–380 ms in Monkey S (Fig. 5F, top), and −160 to 140 ms (Monkey M; Fig. 5E, middle) and −60 to 100 ms (Monkey S; Fig. 5F, middle) relative to saccade onset for saccade direction. Together with the earlier onset latencies in FEF, these results indicate that FEF leads LIP in both the timing and the trial-to-trial dynamics of category and saccade encoding within this reciprocally connected frontoparietal circuit^26^.

We next asked whether the corresponding single-trial latency and flow relationships extend to SC (Fig. 5C–F; purple points in Fig. 5C,D and bottom row in Fig. 5E,F). For category (Fig. 5C,D, left; purple points), FEF preceded SC in both animals, with mean latency difference (SC − FEF) = 83.2 ms in Monkey M (N = 366 trials; p < 0.001) and 39.5 ms in Monkey S (N = 314 trials; p < 0.001). For saccade direction (Fig. 5C,D, right; purple points), the FEF–SC latency relationship was inconsistent between animals: mean (SC-FEF) = 50.1 ms in Monkey M (N = 439 trials; p < 0.001) whereas mean (SC-FEF) = −4.4 ms in Monkey S (N = 331 trials; p = 0.489). Finally, although the onset-latency analysis placed FEF ahead of SC in both monkeys, the directed predictability gain analysis for category (Fig. 5E,F, bottom) showed a clear FEF→SC asymmetry only in Monkey M (significant net FEF→SC flow ∼200–350 ms post-stimulus; p < 0.05, colored dots); in Monkey S the FEF→SC and SC→FEF gains were comparable.

Supplementary analyses showed that saccade-direction directed predictability gain between FEF and SC was not consistent across animals: Monkey M showed stronger FEF→SC around saccade onset, whereas Monkey S showed SC→FEF dominance near onset with only a brief pre-onset FEF→SC bias (Supplementary Fig. S7C,F). For LIP–SC, we did not identify reliable category-related coupling (no clear latency separation and no asymmetry that was consistent across animals), and saccade-related coupling differed across monkeys: Monkey S showed clearer evidence for SC leading LIP, whereas Monkey M showed a more interleaved pattern (Supplementary Fig. S7A,B,D,E). These inconclusive findings do not rule out LIP–SC coupling, because sensitivity may depend on how the decoding axes and resulting task-variable codes are constructed, which we leave for future work.

### FEF inactivation impairs RC task behavior for both spatial orientations of motion and saccade targets

Our population analyses reveal that in both monkeys FEF consistently leads SC and LIP in abstract category encoding, and primarily leads LIP in saccadic-choice encoding. Further, our single-trial analyses support these findings, suggesting that information about category flows from FEF to LIP and SC, and information about saccade plans flows from FEF to LIP. Together, these results suggest that FEF plays an important role in transforming motion evidence into categorical decisions and saccade choices. We causally test this hypothesis by reversibly inactivating right-hemisphere FEF during RC task sessions (Fig. 6A; Table S6). Our group has previously used injections of the GABA_A_ agonist muscimol to demonstrate the causal contributions of both LIP and SC to motion-direction categorization in related tasks, including delayed match-to-category and reaction-time motion categorization paradigms^10,11,20^. Here, we apply this method to reversibly inactivate FEF and causally test its preferential roles in rapid categorical decision-making, as suggested by our neural population analyses of this circuit in the RC task.

**Figure 6.**
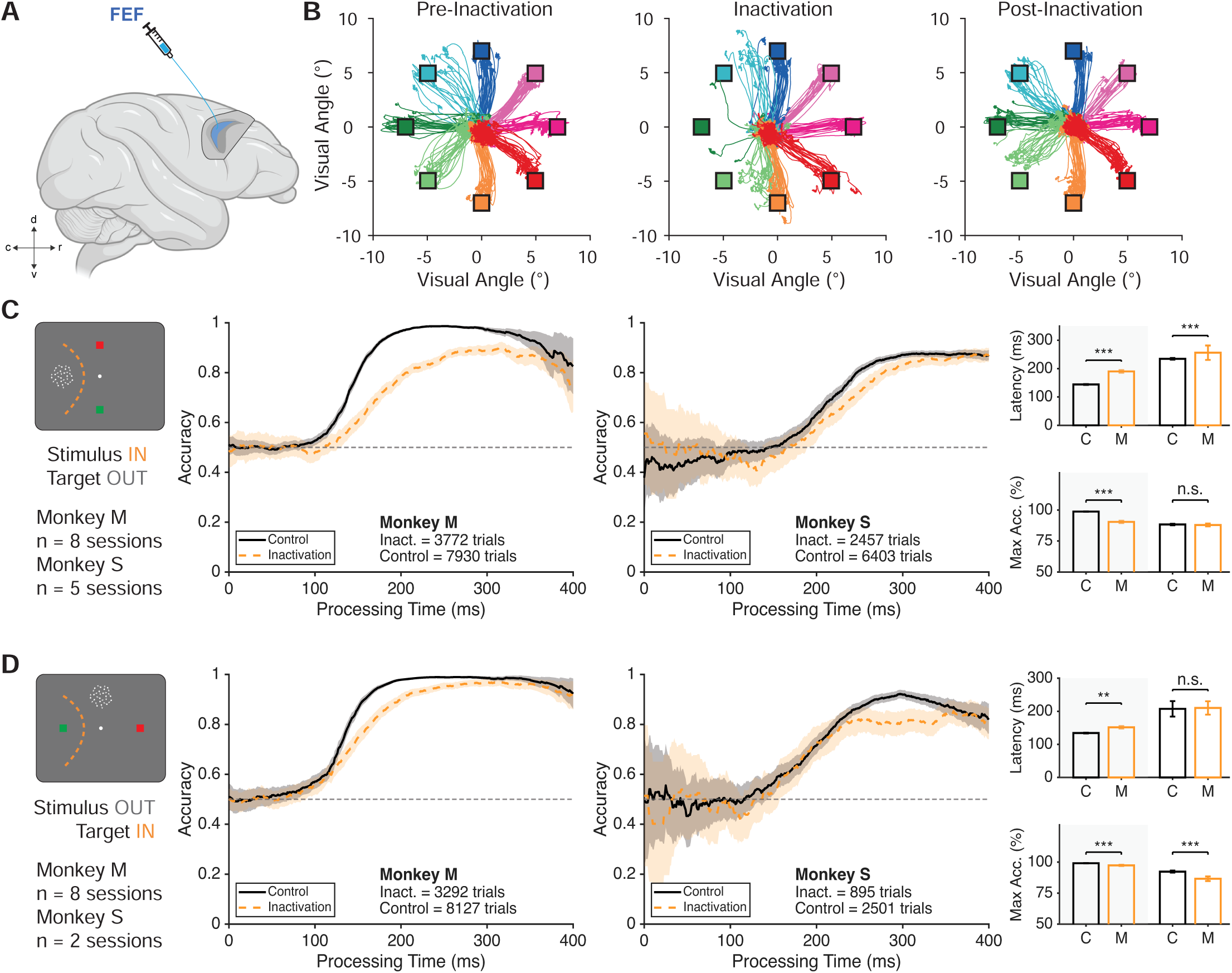
Reversible inactivation of FEF significantly delays and impairs rapid categorization performance. (A), Schematic depicting the targeting of FEF with a GABAA agonist, muscimol. **(B),** Verification of inactivation efficacy with raw gaze traces of correct trials in the memory-guided saccade task. Left panel depicts gaze traces from the day before an inactivation session. Middle panel depicts gaze traces during inactivation; the lack of trials for the leftward target location reflects the lack of correct trials to that location. Right panel depicts the restoration of behavior on the day following an inactivation session. **(C),** Categorization performance in the vertical-target RC task orientation (e.g., motion stimulus positioned within inactivated field, IF) is significantly delayed. Left panel depicts the location of stimulus and targets in this block, with the orange dashed line approximating the IF. Middle panels depict each animal’s tachometric accuracy, in the same format as **Fig 1C**. Dashed orange lines reflects the animal’s accuracy during inactivation. Shaded areas represent 95% confidence intervals calculated using binomial statistics. Right panels show bar plots of the animal’s latency, defined as the time point the animal’s accuracy crossed 0.75 for 150 consecutive time bins. Error bars denote ± one standard deviation across bootstrap iterations. *** denote p<0.001, ** denotes p<0.01, * denotes p<0.05, and n.s. denotes p>0.05 from the pooled-null permutation test. In bar plots, ‘C’ denotes control behavior and ‘M’ denotes behavior after muscimol infusion. Bars on shaded backgrounds depict metrics from Monkey M, while bars on white backgrounds depict metrics from Monkey S. **(D),** The same format as **c**, now quantifying performance in the horizontal-target RC task orientation with a vertically positioned motion stimulus (e.g., one saccade target positioned within IF).

In each inactivation session, we targeted FEF sites where, in prior RC task recording sessions, we observed neural activity which was modulated by the RC task and spatially selective to contralateral stimuli in the MGS task. During inactivation sessions, we verified the efficacy and spatial specificity of inactivation by measuring its impact on MGS behavior (Fig. 6B). Consistent with classic FEF inactivation studies, muscimol produced robust, reversible deficits in memory-guided saccades toward targets in the contralateral visual field—reduced probability of making contraversive saccades, increased errors and hypometric endpoints—while sparing ipsiversive saccades, confirming that our injections produced focal, behaviorally effective inactivation of FEF^27,28^.

To assess how FEF inactivation impacts processing of the motion stimulus and its category, we first examined performance in the vertical-target RC task orientation in which the motion stimulus was presented in the inactivated (left) visual field and the saccade targets were arranged vertically, largely outside the inactivated visual field (Stimulus IN, Target OUT; Fig. 6C). In this arrangement, any deficit should primarily reflect impaired use of motion information rather than a motor impairment for the saccades. Across 8 inactivation sessions in Monkey M and 5 inactivation sessions in Monkey S, we quantified performance as a function of processing time (PT) and computed the latency at which accuracy reached a criterion of 0.75 for 150 consecutive time bins, as well as the maximum accuracy across PTs (see Methods). In both monkeys, FEF inactivation produced a rightward shift of the PT–accuracy curves: the latency to reach criterion was significantly longer during inactivation than in control (Monkey M: 144 ± 1.6ms control vs 190 ± 4.8 inactivation; Monkey S: 234.6 ± 2.9ms control vs 256.2 ± 25.4ms inactivation). In Monkey M, this corresponds to a 47-ms delay (95% CI: 35-55ms; p<0.001 pooled-null permutation test). In Monkey S, this corresponds to a 25-ms delay (95% CI: 11-37.5; p<0.001 pooled-null permutation test). Maximum accuracy was also significantly reduced in Monkey M, from 98.75% in control to 90.4% during inactivation (-8.47% difference; 95% CI [-10.31%,-6.19%; p<0.001 pooled-null permutation test]. In Monkey S, however, maximum accuracy was not significantly reduced, changing from 88.3% in control to 87.9% during inactivation (-0.4% difference, 95% CI [-2.95%, 2.1%]; p=0.278 pooled-null permutation test).

To assess how FEF inactivation impacts saccade planning and execution, we examined performance in the horizontal-target RC task orientation in which the motion stimulus was presented above fixation, outside the inactivated field, while one of the saccade targets was positioned within the inactivated (left) field (Stimulus OUT, Target IN; Fig. 6D). In this geometry, any deficit should primarily reflect impaired target selection, saccade planning, or saccade execution rather than impaired motion processing or categorization. We analyzed 8 inactivation sessions in Monkey M and 2 inactivation sessions in Monkey S with sufficient trials in the horizontal-target block. In Monkey M, FEF inactivation significantly delayed performance: the latency to reach the 0.75 accuracy increased from 134.2 ms in control to 151.5 ms during inactivation, corresponding to a 17-ms empirical delay (95% CI: 10-25 ms; p = 0.010, pooled-null permutation test). Maximum accuracy was also modestly but significantly reduced, from 99.1% in control to 97.39% during inactivation (empirical difference:-1.84%; 95% CI:[-2.87%, - 0.57%]; p < 0.001). In Monkey S, criterion latency was not significantly delayed during inactivation, increasing only from 207.4 ms in control to 210.2 ms during inactivation (empirical difference: 4 ms; 95% CI: [-24 to 29 ms] p = 0.481, pooled-null permutation test). However, maximum accuracy was significantly reduced, from 92.34% in control to 86.59% during inactivation (empirical difference:-6.90%; 95% CI: [-9.64%,-1.43%]; p < 0.001 pooled-null permutation test). Thus, when a saccade target was placed in the inactivated field, FEF inactivation impaired RC task behavior in both animals, but the expression of this impairment differed: Monkey M showed both a modest slowing and a small reduction in asymptotic accuracy, whereas Monkey S showed a selective reduction in maximum accuracy without a reliable latency shift.

As a control for non-specific effects of the injection procedure, we also performed phosphate-buffered saline (PBS) infusions into FEF in Monkey M (2 sessions; Fig. S8; Table S6). In both spatial orientations of the RC task (Stimulus IN/Target OUT and Stimulus OUT/Target IN), PBS infusion did not significantly alter the PT–accuracy curves or choice biases compared to control sessions. Neither the latency to reach the 0.75 accuracy criterion nor the maximum accuracy across PTs differed significantly between PBS and control (bootstrap percentile tests, p > 0.05 for all comparisons). Thus, the delays and impairments in categorization and saccadic choice that we observed are specific to muscimol-mediated inactivation of FEF rather than non-specific consequences of the infusion procedure.

## Discussion

In this study we compared neural population encoding and dynamics across three major nodes of the primate visuospatial and oculomotor network—FEF, LIP, and SC—while monkeys performed an urgent abstract motion categorization task that dissociates stimulus category from the direction of the reporting saccade. Simultaneous recordings showed that neurons in all three areas multiplex motion direction, abstract category, target configuration, and saccade direction. Pseudopopulation decoding revealed a temporal dissociation across areas: LIP and SC provided the earliest representation of task-relevant visual features (motion direction and target identity), whereas FEF showed the shortest-latency encoding of abstract stimulus category. All three areas encoded saccade direction in advance of the monkeys’ decisions, with FEF and SC primarily leading LIP. Leveraging our simultaneous recordings to assess directed interactions across areas at the single-trial level, we found further that category-and choice-related encoding emerges earlier in FEF than LIP and that trial-to-trial fluctuations exhibit directed cross-area predictability consistent with greater information flow from FEF→LIP than the reverse. Results of single-trial analysis also indicated directed flow of category information from FEF to SC. Finally, reversible muscimol inactivation of FEF impaired performance in the RC task, demonstrating a causal contribution of FEF to both accurate categorization and saccadic choice. Together, these results identify a greater contribution of FEF to abstract categorical decisions than expected based on the textbook view of FEF as a core region for oculomotor and attentional control.

Abstract cognition and categorical decision-making have traditionally been linked to canonical executive areas such as LPFC, where earlier work demonstrated robust neural encoding of categories and task rules^29,30,31,32^. However, our categorization studies have revealed that abstract category representations extend well beyond LPFC, with other regions appearing to play a more central role in categorical decisions: parietal area LIP encoded categories more strongly and with a shorter latency than LPFC^14^, while the earlier motion-processing area MT encoded only the sensory features of stimuli rather than their category^8^. This finding was subsequently reinforced when we demonstrated that reversible LIP inactivation impairs categorical decisions in tasks using either saccadic or manual decision reports^10,11^. Extending beyond cortex, we then found that the SC—traditionally linked with reflexive spatial orienting and gaze control—exhibits especially robust, short-latency category encoding, and that SC inactivation produces marked categorization deficits even when decisions are reported with a hand rather than an eye movement^20^. Together, these observations motivated a direct test of where abstract category and saccade-planning signals emerge within the oculomotor circuit. By simultaneously comparing FEF, LIP, and SC in the current study, we identify FEF as the strongest candidate to date for generating categorical choice signals and broadcasting them to LIP and SC–further supported by our reversible inactivation of FEF in the RC task.

A major technical and conceptual advance of the present study is the large-scale simultaneous recording of neuronal populations across FEF, LIP, and SC during decision-making behavior. This approach is especially powerful because it allows us to ask not only where in this circuit category signals first emerge, but also how stimulus representations are transformed into the animal’s categorical decision and ultimately into the eye movement to report the decision. In the present task, stimulus category is dissociated from saccade direction, yet the decision is expressed through a circuit in which FEF and SC are central for oculomotor planning and generation. Simultaneous recordings across all three areas therefore provide a rare opportunity to bridge abstract decision formation and motor implementation within the same trials and the same network. Indeed, the present results already illustrate this advantage. By leveraging simultaneity, our trial-based analyses moved beyond pseudopopulation comparisons and showed that category-and saccade-related signals emerged earlier on single trials in FEF than in LIP, and that trial-to-trial fluctuations were more predictive from FEF to LIP than in the reverse direction. We further found evidence that category-related dynamics in FEF preceded those in SC, linking abstract decision signals to downstream oculomotor structures within the same trials. Overall these findings provide novel insights into directed interactions within the FEF-LIP-SC network, while also motivating future work using even larger-scale simultaneous multi-area population recordings. Applying related analyses to even larger multi-area populations should make it possible to define the geometry of shared and area-specific population codes^33^, identify communication subspaces through which task-relevant information is exchanged^34^, and determine how moment-to-moment interareal dynamics shape reaction times, errors, commitment, biases, and the transformation of decisions into actions.

FEF is classically viewed as a core oculomotor structure involved in selecting and generating saccadic eye movements^35^, and it is also a key node for attentional selection^36,37,38^: subthreshold FEF microstimulation can enhance visual cortical responses and improve covert attention with spatial specificity^39,40,41^. Beyond these roles, visuomotor prefrontal circuits can support deliberation beyond the specification of an action plan; for example, Charlton and Goris showed that population activity first reflects formation of a perceptual choice before transitioning to a motor-plan representation^42^. A close precedent for such abstract categorization in FEF is the Ferrera group’s demonstration that FEF neurons track category boundaries during slow–fast motion-speed categorization and shift with changes in decision criteria^43^. Our work builds on this foundation by revealing that category and choice signals emerge earliest in FEF among the oculomotor triad, underscoring how strongly this network (and FEF in particular) can be engaged by abstract, rule-defined decisions.

A central mechanistic question is raised by these findings: what is the coding scheme by which the FEF neural population simultaneously encodes both abstract category information and the impending saccadic choice? Many single neurons are modulated by both variables, arguing against a strict division into “cognitive” versus “motor” cell classes. Instead, the mixed selectivity for category and saccade-choice suggest that these signals occupy separable representational subspaces in FEF, related to similar results we previously reported for SC^20^. This representational geometry offers a concrete mechanism for multiplexing: FEF can support abstract, rule-based categorization while simultaneously participating in action planning, with downstream circuits selectively reading out categorical versus choice information from different population dimensions. Notably, the same organizing principle emerged in our prior SC study, suggesting that subspace separation may be a shared strategy across oculomotor circuits for combining–and potentially linking– abstract decision variables with motor-related signals.

The current study replicates and extends our previous surprising finding that the SC is meaningfully engaged in abstract visual categorization. A key reason this result was unexpected is that prior to that SC study, LIP had been our leading candidate brain area for mediating categorical decisions in our motion categorization framework^1^. Against that backdrop, and reinforced by the present results, Peysakhovich et al. uncovered not only that category signals in SC emerge reliably and with shorter latency than those in LIP, but also that SC perturbation produces striking categorization deficits even when decisions are reported with a hand rather than an eye movement—demonstrating that SC’s involvement is not a trivial byproduct of saccade preparation. In the present urgent, saccade-report paradigm, category encoding emerges even earlier in FEF than SC, and our cross-area analyses indicate that SC category encoding is consistent with driving input from FEF. Together, these findings support a framework in which early categorical and choice signals in FEF rapidly project to SC, positioning SC to influence the timing and expression of the emerging choice within oculomotor selection circuitry, consistent with prior work implicating SC in the transition from deliberation to action^44^.

The compelled-saccade “urgent choice” framework developed by Stanford, Salinas, and colleagues offers a powerful and principled way to relate neural dynamics to the window of evidence available for decision formation^,21,22,45^. Relative to our more commonly used delay-based categorization paradigms, which are powerful for isolating stimulus processing and response planning but necessarily engage recurrent short-term memory and comparison processes, the urgency manipulation in the present task constrains interpretation to a narrow temporal epoch in which sensory evidence can influence choices. By aligning analyses to processing time (and the resulting tachometric function), urgent-choice designs help separate neural signals associated with early evaluation of stimulus evidence from later commitment and movement execution, and they make cross-area latency comparisons cleaner by controlling when informative evidence becomes available to the animal. This is particularly useful in abstract categorization, where the core computation is the transformation of sensory evidence into a learned, rule-defined meaning rather than veridical readout of motion direction. A key future direction is to use urgency to estimate when category representations become “decision-effective” (i.e., predictive of commitment) and whether that threshold is set within FEF or emerges from cooperative interactions within the FEF-LIP-SC circuit, and to test these inferences with temporally precise perturbations paired with simultaneous multi-area recordings.

An important question is how broadly these oculomotor-circuit contributions to abstract categorization generalize across task designs, response modalities, and sensory domains. The present results show that robust category signals which we previously observed in SC and LIP during a delayed match to category paradigm with a manual decision report^20^ extend to urgent decisions reported by saccade. This indicates that oculomotor-network engagement is not restricted to particular paradigms with extended processing windows or delay-period demands. It is also consistent with other work from our group showing that LIP category encoding generalizes between tasks requiring saccadic and manual responses^46^. We hypothesize that we would find a similar pattern of results, with FEF playing a preferential role relative to SC and LIP, if we tested a categorical decision task in which decisions were not reported via eye movements. However this will need to be explicitly tested in future work. A related question is whether the recruitment of the FEF–LIP–SC circuit is specific to motion categorization because of its close links to dorsal-stream processing, including MT^47^. We think this is unlikely: LIP category signals generalize beyond motion, including to learned associations among non-motion stimuli such as shapes^48^, supporting the view that these signals reflect learned associative or semantic relationships rather than a motion-specific processing. Together, these observations motivate future tests of generality across stimulus domains, sensory modalities, and behavioral paradigms, including more naturalistic reward-based foraging tasks^49^ that monkeys can rapidly learn to perform—an important complement to well-controlled paradigms given growing evidence that training history can shape category representations and circuit engagement^12,50^.

Going forward, we strive to understand why the FEF-LIP-SC network–traditionally viewed as fundamentally anchored in spatial processing, attention, and gaze control–is recruited to mediate non-spatial abstract cognition. We recently identified a robust behavioral signature of the engagement of this network by abstract categorical decisions: small-amplitude (< 1.0 degree) but reliable category-correlated gaze shifts, particularly during the working memory period of the delayed match to category task.^51^ An intriguing speculation is that this network’s spatial axes for representing the external world might be repurposed in order to represent and manipulate non-spatial abstract cognitive variables–akin to using spatial strategies to solve non-spatial problems such as using a number line to solve an arithmetic problem or visualizing data on a Cartesian plot.

## Author contributions

Conceptualization: DJF, OZ, VS. Experiments: OZ, VS, SD. Formal analysis: OZ, MG, YX, VS. Visualization: OZ, MG, YX. Writing – DF, OZ, MG. Writing – editing: OZ, VS, SD, MG, YX, BD, DJF. Funding acquisition: DJF and BD. Project supervision: DJF.

## Acknowledgments

We are grateful for helpful discussions and experimental input from Christopher Hauser and expert assistance from the staff of the UChicago Animal Resources Center. We thank Rory Cooley and Matthew Rosen for input on an earlier draft of this manuscript. This work was supported by NIH R01EY019041 (DJF), NIH R01EY037119 (DJF and BD), DOD Vannevar Bush Faculty Fellowship N000141912001 (DJF), and grants from the NSF (DMS-2235451) and Simons Foundation (MPS-NITMB-00005320) to the NSF-Simons National Institute for Theory and Mathematics in Biology (NITMB), and NIH 1F30EY033648 (OZ).

## Data Availability

The data analyzed in this study will be posted to FigShare (www.figshare.com) at the time of publication.

## Code Availability

Data analysis code developed and used to generate these results will be made available at the time of formal publication.

## Methods

### Subjects

Two adult male rhesus macaques (*Macaca mulatta)* participated in the experiments: Monkey M (21-24 years old; ∼10kg) and Monkey S (9-12 years old, ∼13 kg). All procedures were in accordance with the University of Chicago Institutional Animal Care and Use Committee and the National Institutes of Health guidelines and policies.

### Behavioral tasks

For all behavioral tasks, the monkeys were head restrained and seated in a primate chair that was either inserted inside an isolation chamber (Crist Instruments) or positioned in a dedicated experimental room. Visual stimuli were presented on a 21-inch color CRT monitor with a resolution of 1280×1024 at 75 Hz refresh rate, viewed from 57 cm. Gaze positions were tracked using an Eyelink 1000 optical eye tracker (SR Research) at a sampling rate of 1 kHz. Task events, stimulus presentation, behavioral signals, and reward delivery were controlled by the MATLAB-based toolbox NIMH MonkeyLogic^52,53^ on a Windows-based PC.

### Memory-Guided Saccade Task

To identify visual and motor fields of LIP, FEF, and SC neurons, we trained monkeys to perform a memory-guided saccade (MGS) task. For this task, monkeys initiated trials by acquiring and maintaining fixation on a central white fixation point (0.2° radius) within a 2.5° radius fixation window. After 500ms of fixation, a 0.6° white square target was briefly presented at one of eight peripheral locations at 7° eccentricity for 300ms. After a 1000 ms delay period, the fixation point disappeared and signaled the monkeys to make a saccade to the remembered target location to receive a juice reward. Recording coordinates in all brain areas were selected based on the presence of neurons with visual or motor fields in the left visual field, which was contralateral to the recorded hemisphere. In every recording session, around 100 trials of the MGS task were first completed prior to starting the rapid categorization task.

### Rapid Categorization (RC) Task

The RC task was designed to compel the animals’ to make rapid decisions about the abstract category membership of visual motion stimuli, and to report those decisions by their saccadic eye movements. All three of the brain areas targeted for recordings are known to contain neurons which encode visual stimulus features of the stimulus in their RFs, in addition to encoding aspects of the monkeys’ current and planned eye movements. Neurons also can show mixed-selectivity, which manifests as encoding of multiple stimulus, motor, and task-related factors. In order to assess and compare neurons’ visual stimulus encoding and saccade-related encoding, we tested the RC task in two blocks (300-500 correct trials per block) which tested different geometries of visual stimulus and target locations. In the vertical-target orientation of the RC task, the visual motion stimulus was shown 7° to the left of the fixation point, and the saccade targets were 7° above and below fixation. Since we were recording from the right hemisphere, this places the motion stimulus in a location likely to stimulate many neurons with RFs overlapping the contralateral hemifield (Fig. S3). In the horizontal-target orientation of the task, the motion stimulus was positioned 7° above the fixation point and the saccade targets were positioned 7° left and right of the fixation point. This spatial orientation is designed to examine neural activity related to saccade plans toward vs away from the contralateral visual field. Each recording session comprised 300-500 completed trials for both spatial orientation of the RC task.

In contrast to previous motion categorization studies from our group, the random-dot stimulus shown at the start of each RC task trial was a 100% noise stimulus (i.e. 0% motion coherence), and switched to a 100% coherent stimulus later in the trial. This noise stimulus occupied the same size and position as the informative motion stimulus that appeared later in the trial. The purpose of this noise stimulus was to minimize visual onset transients in neural activity and bottom-up attention capture when the informative stimulus was shown.The 100% coherent motion stimuli consisted of 5° diameter circular patches of ∼100 high-contrast dots moving at 12°/s. Six directions spanning 360° were divided into two arbitrary categories that were each arbitrarily paired with either a red or green saccade target (Red: 75°, 135°, 195°; Green: 255°, 315°, 15°).

Monkeys initiated trials by acquiring and maintaining fixation on a central fixation point (0.2° radius) within a 2.5° radius fixation window. After 500 ms of fixation, the 0% coherence random dot motion stimulus (5° diameter) appeared at the appropriate location on the display depending on whether it was a vertical-target or horizontal-target orientation block. After 300ms of continued fixation, red and green targets (1° square) appeared at either 7° above/below or left/right of the fixation point for vertical-target or horizontal-target orientation blocks, respectively, with their relative positions randomized on each trial to decouple saccade-target locations (and the monkeys’ saccade directions) from motion categories. After continued fixation for a randomized target time, the fixation point disappeared, which served as the ‘go cue’ for the monkeys to initiate a saccade within 500ms. For Monkey M, target times were sampled uniformly from 250, 350, and 450ms; for Monkey S, target times were sampled from 400ms and 500ms at a 2:1 ratio, respectively. After a gap time which followed the go cue and was randomized on each trial, the noise stimulus was replaced by an informative motion stimulus (with 100% coherence). For Monkey M, gap times were sampled uniformly from [0, 15, 50, 90, 130, 175] ms; for Monkey S, gap times were sampled from [-100, 0, 100] at roughly a 2:1:1 ratio. Gap times ranged from 0-250ms for Monkey M and-100-100ms for Monkey S; these gap time ranges for each monkey were chosen to target an overall accuracy of 80% to maintain a level of correct performance in order to sustain a high level of motivation in the subjects. Monkeys received a juice reward for making a saccade to the correct color target and maintaining fixation on the target for 300ms.

### Surgical procedures and electrophysiological recordings

Prior to behavioral training, we obtained an MRI scan of each monkey’s head. Then, monkeys were implanted with a titanium headpost and, after they were trained on both tasks, two MRI-guided, custom titanium recording chambers were implanted above their right hemisphere (Rogue Research). One chamber was positioned over the frontal eye fields (FEF), and the other was positioned over LIP and SC. Recordings were conducted acutely using a combination of 24-or 32-channel dense linear arrays (Plexon V-probes or S-probes). Up to six probes were used within a session to simultaneously record from LIP, FEF, and SC coordinates with two probes per area (up to four probes within one recording chamber; see Fig. 2B). Probes were lowered into dura-piercing guide tubes and positioned into each area using NAN microdrives (NAN instruments). Neurophysiological signals were amplified, digitized, and stored using the Plexon Omniplex acquisition system. We used anatomical landmarks and responses during the MGS task to confirm the targeting of each probe in the correct brain area. For SC recordings, we primarily targeted neurons in the intermediate layers, although other layers were often sampled due to the ∼2mm span of recording channels on the probes. Spike sorting was performed with the MATLAB-based Kilosort 2 toolbox, which was run on a Windows-based PC equipped with an NVIDIA RTX 3070 graphics card. Putative neuron clusters identified by Kilosort 2 were further curated using the Python toolbox Phy. Well isolated single units were distinguished from multi-unit activity using criteria that includes: high amplitude template waveforms, waveforms isolated from all other putative units in at least one channel across the length of the recording, and <1% of spikes with an inter-spike interval of <2ms. Coordinates in which we recorded FEF were distinguished from pre-frontal cortex using low amplitude microstimulation, such that saccades were able to be elicited with 200ms of biphasic pulses as low as 50 µA delivered at 200 Hz.

### Reversible inactivation experiments in FEF

We adapted the protocol developed by Vanegas et al. 2019 to inactivate FEF coordinates^54^. We infused muscimol, a GABA_A_ agonist (typically 1.5-2 µL of 5 µg/µL solution), into previously recorded FEF coordinates with response fields to the contralateral left visual hemifield. The details regarding each inactivation session are tabulated in Table S8. The muscimol solution was prepared by dissolving solid muscimol (Sigma Inc.) in phosphate-buffered saline (PBS). To ensure accurate targeting of grey matter for inactivation, we monitored neural activity prior to infusion using either custom-made injectrodes (Monkey M) or 16-channel Plexon S-probes with an embedded fluid cannula (Monkey S). Thirty minutes post-infusion, monkeys performed a block of the MGS task to assess and confirm behavioral deficits from inactivation (Fig. 6B), followed by two blocks of the RC task—one for each target orientation. Due to the large number of trials required for a complete dataset of the RC task, we were unable to obtain pre-and post-inactivation data within the same session. Instead, we used behavioral data from the day before and the day after each inactivation session as controls to compare to behavior during inactivation. In total, we analyzed data from 13 inactivation sessions (Monkey M: 8 sessions, Monkey S: 5 sessions), and 2 saline control sessions (Monkey M). During the process of performing inactivation experiments in Monkey S, the animal experienced unrelated health and behavioral complications which precluded any further experiments.

### Data analysis

Analyses were performed using MATLAB (R2021a-R2025b). For the RC task, behavioral analyses focused exclusively on ‘completed trials’ where the animal successfully made a saccade to a color target after the go cue, including both correct and incorrect trials. On each trial, the onset time of the saccade was determined using gaze-tracking data, which was smoothed using a 3ms moving average filter and then differentiated to estimate gaze velocity. Saccade onset was identified as the moment when gaze velocity was last under 50°/s prior to the peak saccade velocity. Processing time (PT) was calculated for each trial as the duration between the time the 0% coherence noise stimulus was replaced by the 100% coherence motion stimulus, and the time of saccade onset. For neural analyses, unless otherwise noted, spike trains were smoothed using a Gaussian kernel (σ=20ms).

### Behavioral analyses

To assess how the animal’s behavioral performance evolved as a function of processing time (PT), we pooled all completed trials from each animal across all recording sessions and computed tachometric curves separately for each monkey and stimulus-target spatial configuration. Trials lacking a valid saccade to a color target from the animal were excluded. Correct trials were defined as trials in which the animal made a saccade to and fixated on the correct color target for 300ms. Incorrect trials were defined as trials in which the animal made a saccade to an incorrect color target or broke fixation from either color target within 300ms. Tachometric curves were evaluated across a PT range of 0–400 ms in 1 ms time steps. At each PT center t, we included trials with PT ∈ [*t*−40,*t*+40], yielding an 81 ms sliding window.

To reduce the influence of local category imbalances, especially at low PT ranges (<100ms) where trial counts were fewer, accuracy was first computed within each category before averaging across categories. For category *c* at PT center *t*, let *n_c_*_,*t*_ denote the number of trials in the local PT window and *k_c_*_,*t*_ denote the number of correct trials. The category-specific accuracy was estimated using the Agresti–Coull method^55,56^:

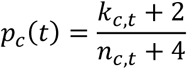

When both categories were present in the window, the category-balanced empirical accuracy was defined as the equal-category average:

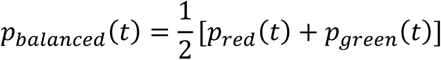

If only one category was present in a sparse early or late window, the available category estimate was used; if no trials were present, the value was left undefined. This method estimates the expected value of the animal’s accuracy as if each category had the same number of trials without stochastic subsampling.

Pointwise 95% confidence intervals were computed from the same category-balanced estimator. For each category, the stabilized binomial standard error was

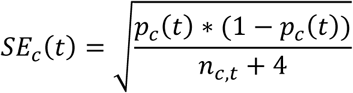

When both categories were present, the category-balanced standard error was

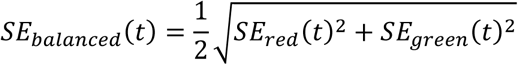

and the 95% confidence interval was computed as

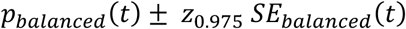

When only one category was present, the interval was computed from that category’s stabilized binomial standard error. Confidence intervals were clipped to the valid accuracy range of 0 to 1. Tachometric curves plotted in figures show the empirical category-balanced tachometric estimate, with a pointwise 95% confidence band and a horizontal reference line at chance performance of 0.5.

### Quantifying feature selectivity of single neurons

The firing rate of a neuron on single trials may vary depending on several features, which include the direction of the motion stimulus, the category of the motion stimulus, the relative positions of the saccade targets (e.g. red target above, green target below), or the animal’s saccade direction. A single neuron may be selective to all these features simultaneously, so we fit a nested ANOVA model to the neuron’s firing rates at four time points relative to the category stimulus onset: (100ms, 200ms, 300ms, and 400ms). This analysis only includes correct trials on which the animal had 200ms or greater processing time, and spike trains were smoothed using a Gaussian kernel (σ=40ms). For ANOVA factors, we used the stimulus category (red = 1; green =-1), the motion direction nested within category, the saccade direction (up = 1; down = - 1), and a target configuration regressor which was computed as an interaction term of category x saccade. Trials corresponding to different task orientations (in which the stimulus was positioned above the fixation point rather than in the left visual field) were analyzed separately, and a neuron was considered selective for each of these factors if its minimum p-value across the four time-points met the criterion in either task orientation (Bonferri-corrected α = 0.05/8).

### Visualizing single neuron firing rate modulations by feature

To visualize neuronal activity, we aligned each neuron’s firing-rate time series to one of three event times: (i) saccade-target onset, (ii) onset of the 100% coherence motion stimulus, and (iii) saccade onset. For each alignment, firing rates were averaged across trials grouped by the feature most relevant to that epoch: for saccade-target onset, trials were grouped by the position of the red target (up vs. down); for motion-stimulus onset, by the six motion directions; and for saccade onset, by saccade direction. For Fig. 2, visualization was restricted to neurons that were significantly selective for all of these features, as assessed by ANOVA.

### Receptive Field Mapping

To determine the visual receptive fields and the movement response fields of feature-modulated neurons and multi-unit clusters, we analyzed neural activity during each of the eight peripheral target locations of the MGS task. To capture a range of FEF, LIP, and SC neural MGS response profiles, activity from 0 to 300 ms following visual target onset was used to assess visual receptive fields, and activity from 300 to 0 ms preceding saccade onset was used to determine movement response fields. Units with activity significantly modulated by MGS target location (one-way ANOVA, p < 0.01) during the analyzed time windows were included in each analysis, respectively. A spike count by condition direction vector sum was then used to determine each unit’s preferred visual or saccade location, with the distributions of each area and each monkey visualized in a polar histogram (Fig. S3).

### Quantifying single-neuron category tuning

In prior work, we have used a receiver operating characteristic (ROC)-based category tuning index (rCTI) to distinguish category selectivity from direction tuning in single neurons. This analysis was restricted to correct trials on which the animal had at least 200 ms of processing time. To quantify the magnitude and time course of category selectivity in the RC task, we adapted that approach to the specific set of motion directions in our experiment. For each neuron previously identified as category or direction selective (by nested ANOVA), we aligned their smoothed firing rate to the onset of the 100% coherence motion stimulus. At each time bin (1-ms step), we formed trial-by-trial firing-rate distributions for each motion direction and quantified discriminability between pairs of directions using the area under the ROC curve. For each direction pair, the direction with the larger mean firing rate was assigned as the positive class and discriminability was quantified as the rectified AUC (0.5 + |0.5-AUC|). We computed rectified AUCs for within-category (WC) direction pairs (e.g. 75° and 135°) and between-category (BC) pairs (e.g. 75° and 15°). To balance the number of direction pairs between and within categories spaced 60° and 120° apart, we weigh the 120° WC pairs (75° and 195°, 15° and 255°), and the 60° BC pairs (75° and 15°, 255° and 195°) twice in calculation. This produces four pairs spaced 60° apart and four pairs spaced 120° apart for both WC and BC. We excluded direction pairs separated by 180° as all pairs would be in different categories. The rCTI at each time bin was defined as the mean rectified BC AUC minus the mean rectified WC AUC:

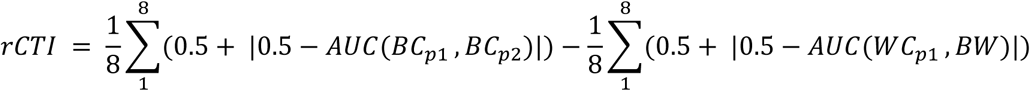

where BC_p1_ and BC_p2_ are the two directions in the p^th^ BC pair, and WC_p1_ and WC_p2_ are the two directions in the p^th^ WC pair. Population rCTI time courses were obtained by averaging rCTI time courses across neurons, and error bounds were quantified as SEM across neurons.

### Pseudopopulation support vector machine analysis

To quantify the magnitude and temporal evolution of population-level selectivity within each brain area, we trained linear support vector machine (SVM) decoders to predict task features from firing rates of pseudopopulations assembled by pooling units across recording sessions. Units used for pseudopopulations included both single neurons and multi-unit clusters. To match population sizes across brain areas, we subsampled units without replacement from areas with larger unit counts until each area contained the same number of units as the area with the smallest available population of units.

To ensure that each unit contributed activity from the appropriate behavioral and task context, we first enumerated the full set of unique task-condition combinations, defined by (i) task/target orientation, (ii) motion direction, (iii) saccade direction, and (iv) trial outcome (correct/incorrect). For each unique condition, we generated a fixed number of pseudotrials (e.g., 100). For each unit, we identified all candidate real trials from that unit’s recording session that matched the pseudotrial’s condition and then split those trials into disjoint training and testing pools (80%/20%). Real trials were sampled without replacement when sufficient trials were available and with replacement otherwise. This procedure yielded pseudopopulation activity by combining independently sampled single-trial responses across units, while ensuring that decoder performance was evaluated on test data that were not used during training. Firing rates were aligned to a common event time (e.g., motion stimulus onset or saccade onset), and SVMs were trained and tested on firing rates at a specific time point every 5 ms. Pseudopopulations were reconstructed independently on each run (1000 runs total), and decoder accuracy was averaged across runs, with variability reported as ± 1 SD.

To quantify direction encoding in a category-independent manner, we trained and tested the SVM using motion directions from only one category at a time, separately for each category. Firing rates were aligned to the onset of the 100% coherence motion stimulus, and SVMs were trained and tested at 5 ms time steps. To mitigate effects of category preference in single-unit responses (e.g., systematically lower firing rates for one category), the category yielding the higher decoding accuracy was used for subsequent visualization and latency analyses. To quantify category encoding independent of motion direction, we trained two SVMs to predict category, each using two motion-direction pairs separated by 180°: (75° and 255°, 135° and 315°) and (195° and 15°, 135° and 315°), respectively. We then tested each trained decoder on the one remaining motion-direction pair separated by 180°: (195° and 15°), and (75° and 255°), respectively. Using a simulated population of purely direction-tuned neurons (cosine tuning), we verified that this decoding scheme yields chance-level category accuracy (0.5) for the categories used here. For subsequent visualization and latency analyses, we averaged decoder accuracy across the two training-pair configurations. To decode target configuration or saccade direction, firing rates were aligned to target onset or saccade onset, respectively. Trials from the vertical-target orientation block were used for direction and category decoding, which best positioned the motion stimulus within the receptive fields of most recorded neurons. In contrast, trials from the horizontal-target orientation block, in which one saccade target was placed in the receptive field of recorded neurons, were used for target configuration and saccade-direction decoding.

### SVM latency comparisons

To compare feature selectivity latencies across brain areas, we defined the selectivity latency for each pseudopopulation as the earliest time point at which SVM decoding accuracy exceeded a dynamic performance threshold for five consecutive time bins. The dynamic threshold was set as halfway between chance performance for that decoder and the maximum accuracy attained by the decoder for a given pseudopopulation. For direction-decoding SVMs, the chance performance level is 0.33, and for all other SVMs the chance performance level is 0.5. The minimum allowable threshold for a pseudopopulation was set at 0.1 above chance; for motion direction decoding the threshold floor was thus 0.43, and for all others the floor was 0.6. The windows used for testing latency are as follows: 0 to 400ms from stimulus onset (category and motion direction decoders), 0 to 400ms from target onset (target configuration decoders), and - 300 to 100ms surrounding saccade onset (saccade direction decoders).

To test whether latency distributions differed between two areas, we first computed the observed difference in mean latencies. Statistical significance was assessed using a two-sided subsampled permutation test with 20,000 iterations. On each iteration, the two latency vectors were pooled and randomly subsampled without replacement into two samples of 50 latencies. Then, for each iteration we computed the difference between the means of these samples to form a null distribution. The p-value was estimated as the proportion of permutations for which the permuted difference in latencies equaled or exceeded the observed difference in latencies.

We examined the robustness of our latency measurements in each brain area as a function of neural pseudopopulation size, to test whether the number of units sampled in our experiments affected our results. We repeated the decoding and latency analyses by subsampling units across population sizes ranging from 50 to the minimum available number of units across the three areas, with a step size of 1. As above, the dynamic threshold for each pseudopopulation decoding run was set as halfway between chance performance and the maximum accuracy attained. The minimum allowable threshold for a pseudopopulation was set at 0.1 above chance; for motion direction the threshold floor was thus 0.43, and for all others the floor was 0.6. 100 SVM runs were performed at each pseudopopulation size (but 200 runs for stimulus category decoding: 100 per set of directions; see *Pseudopopulation support vector machine analysis*, above), and mean latency was plotted against pseudopopulation size (Fig. S5).

### Analyses of behavior from inactivation sessions

To quantify the effect of FEF inactivation on the speed and accuracy of behavior, we used a procedure adapted from our main behavioral analyses, described earlier. Analyses were performed separately for each monkey and stimulus-target spatial configuration. Trials lacking a valid saccade to a color target from the animal were excluded. In brief, tachometric curves were computed over a 0-400ms PT range using 1ms steps and an 81ms sliding window. The animal’s accuracy at every window was estimated separately by category with the Agresti-Coull correction and then averaged to correct for trial imbalances between categories.

We then quantified two behavioral metrics from the empirical control and inactivation tachometric curves. First, we estimated a “behavioral latency” as the earliest processing time at which the animal’s accuracy exceeded 0.75 for 150 consecutive time steps. We then computed the latency shift due to inactivation as the latency of inactivation behavior minus the latency of control behavior. Second, we quantified the maximum accuracy the animal attained across the PT range, and we computed the accuracy deficit as the maximum accuracy of inactivation behavior minus control behavior.

To test the significance of these measurements, we generated a null distribution by permuting the ‘control’ and ‘inactivation’ labels of trials to generate two tachometric curves, preserving the original number of trials in control and inactivation datasets, and quantifying their difference in latencies and maximum accuracies across 10,000 iterations. For latency, a one-sided p-value was computed to test the null hypothesis of whether latency was significantly higher for inactivation behavior. P-values were defined as the fraction of null latency differences greater than or equal to the observed latency shift. For maximum accuracy, a one-sided p-value was computed to test the null hypothesis of whether maximum accuracy was significantly lower for inactivation behavior. P-values were defined as the fraction of null maximum accuracy differences less than or equal to the observed accuracy change, testing whether inactivation reduced performance.

### Trial-resolved feature evidence and cross-area flow

To relate population coding to trial-by-trial interactions between simultaneously recorded areas, we performed a two-stage analysis. First, for each session and area we learned a session-wise linear axis for a task feature and used the projection onto this axis as a time-resolved feature evidence signal. Second, we used these evidence traces to (i) define single-trial onset latencies by a sustained threshold-crossing rule (Fig. 5C,D; Fig. S7A,D) and (ii) quantify directed interactions between areas via lagged ridge regression with permutation-based significance testing (Fig. 5E,F; Fig. S7B,C,E,F).

### Spike-count binning, alignment, and within-session normalization

For each session and area, spikes were binned into a tensor aligned to task events using a 20 ms sliding window with a 10 ms step, yielding effective 10 ms temporal resolution. For stimulus-aligned analyses (category; vertical-target trials), the analysis window spanned [−250, 800] ms relative to stimulus onset. For saccade-aligned analyses (saccade direction; horizontal-target trials), the analysis window spanned [−400, 300] ms relative to saccade onset. Time axes refer to window centers. Let **X** ∈ ℝ^L×T×N^ denote binned spike counts, where L is the number of trials, T is the number of time bins, and N is the number of simultaneously recorded units. We normalized each unit within session/alignment by z-scoring across all trials and time bins:

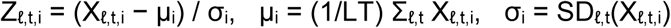

yielding **Z**, which was used for all subsequent axis, onset, and flow analyses. Each trial carried labels for category C ∈ {−1, +1}, motion direction R, saccade direction S ∈ {−1, +1}, target-layout orientation (vertical vs. horizontal), processing time (PT; ms), and correctness. Unless otherwise specified, analyses were restricted to trials with PT ≥ 200 ms and included both correct and incorrect trials. Including all completed trials estimates interareal predictability over the full post-filtered task distribution, whereas restricting the analysis to correct trials would condition the estimate on successful behavior and could remove trial-to-trial variability relevant to the coupling estimate.

### Session-wise feature axes and evidence traces

For each session and area, we learned a linear feature axis and computed a time-resolved evidence trace by projecting population activity onto that axis. Let **z**_ℓ_(t) ∈ ℝ^N^ denote the population vector from **Z** for trial ℓ at time bin t. For a feature F ∈ {C, S}, we learned a unit-norm axis **a**_F_ ∈ ℝ^N^ and defined the scalar feature code:

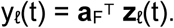

Rather than using fixed training windows, we performed a grid search over candidate windows to identify the optimal window for axis training in each session and area. Candidate windows were generated by varying the window start in 20 ms steps across a search range and testing window durations of 50, 100, 150, 200, and 250 ms. For stimulus-aligned category feature analysis, the search range was [0, 500] ms after stimulus onset. For saccade-aligned saccade direction feature analysis, the search range was [−200, 50] ms relative to saccade onset. For each candidate window, we averaged **Z** within the window to obtain a trial-by-unit matrix and fit an L_2_-regularized logistic regression decoder with class balancing. The inverse regularization strength C was selected by 5-fold stratified cross-validation over C ∈ {0.1, 0.3, 1, 3, 10}. To score each candidate window, we computed out-of-fold per-bin AUC: for each fold, we trained the axis on window-averaged activity from the training set, then evaluated decoding performance at each time bin within the window using test-set trials. The optimal window was defined as the candidate yielding the highest peak per-bin AUC (the maximum AUC across bins within that window), and the final axis was trained on all included trials using activity from that window.

The feature axis was defined as the unit-norm coefficient vector **a**_F_ = **w**_F_ / ‖**w**_F_‖, with sign chosen such that the resulting projection yields AUC ≥ 0.5. For saccade direction axes, we additionally used per-trial sample weights to balance trials across joint (C, S) strata (weights proportional to the inverse stratum count), preventing saccade decoding from being driven by category imbalance. To reduce category contamination in saccade evidence, we constructed an “invariant” saccade axis by orthogonalizing the raw saccade axis to a saccade-aligned category axis (trained on the optimal window within [−300, −50] ms):

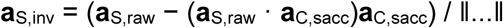

and used **a**_S,inv_ for saccade analyses when available.

*Time-resolved QC and session inclusion*

We evaluated axis quality by computing time-resolved decoding performance from a fixed axis. For each time bin t, we computed the projection p_ℓ_(t) = **a**_F_^⊤^**z**_ℓ_(t) and then computed ROC AUC across trials (Fig. 5A,B). For inclusion in the onset and flow analyses, a session was considered to have a usable feature axis if its QC curve reached AUC ≥ 0.65 at any time bin. For each area pair, we applied symmetric inclusion: a session contributed only if both areas passed the QC criterion for the relevant feature, ensuring identical session sets for both directions in the flow analysis.

### Single-trial feature onset latency

To estimate trial-wise onset latencies (Fig. 5C,D; Fig. S7A,D), we formed a signed evidence trace by multiplying the 1D projection by the trial label sign:

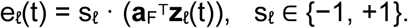

where s_ℓ_ is the ground-truth label for trial ℓ. Evidence traces were smoothed in time with a Gaussian kernel of σ = 20 ms. For each trial, we computed baseline statistics from a pre-event baseline window and defined a trial-specific threshold θ_ℓ_ = μ_ℓ,0_ + 4σ_ℓ,0_, where (μ_ℓ,0_, σ_ℓ,0_) are the mean and standard deviation of e_ℓ_(t) within the baseline window. Baseline and search windows were feature/alignment-specific: for category (stimulus-aligned), baseline [-200, 0] ms and search [0, 500] ms; for saccade direction (saccade-aligned), baseline [-350,-200] ms and search [-300, 200] ms. We defined onset latency as the first time bin within the search window at which e_ℓ_(t) > θ_ℓ_ for at least 5 consecutive bins. Trials without a sustained crossing were assigned no onset. For each area pair, we included only trials with defined onsets in both areas. To assess latency differences, we computed paired trial-wise latency differences and assessed significance with a two-sided sign-flip permutation test on the mean difference (20,000 permutations).

### Directed flow on projected feature codes

To quantify directed interactions in trial-to-trial fluctuations of the feature evidence (Fig. 5E,F; Fig. S7B,C,E,F), we applied a lagged ridge-regression framework to the projected traces. For an ordered pair of areas A → B, let yℓ^A^(t) and yℓ^B^(t) denote the feature projections for trial ℓ at time t in areas A and B, respectively. To remove the session-mean evoked component in the projected space, we subtracted the across-trial mean code at each time bin after smoothing with a Gaussian filter of σ = 10 ms.

We constructed lagged predictors using a lag window specified per area pair and alignment: 80 ms for FEF–LIP and LIP–SC stimulus-aligned (category) analyses (W = 8 bins), 50 ms for FEF–LIP saccade-aligned analyses (W = 5 bins), and 30 ms for FEF–SC (both alignments) and LIP–SC saccade-aligned analyses (W = 3 bins), at the 10 ms effective bin spacing. At each time bin t ≥ W, we compared two ridge regression models predicting the target-area evidence yℓ^B^(t) across trials: (i) a reduced model containing an intercept and the target area’s own history {yℓ^B^(t-1),…,yℓ^B^(t-w)} (autoregressive control), and (ii) a full model that additionally included the source-area history {yℓ^A^(t-1),…,yℓ^A^(t-w)}. Ridge regression used a fixed penalty λ = 10^−2^, and predictors were used without additional standardization beyond the initial within-session z-scoring of neural activity. We quantified directed influence as the log-likelihood improvement from adding the source history, expressed in bits:

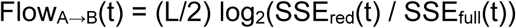

where L is the number of trials and SSE_red_(t) and SSE_full_(t) are sums of squared prediction errors for the reduced and full models, respectively. This metric, which we term “directed predictability gain,” quantifies how many bits of information the source area’s history provides about the target area’s current state beyond its own autoregressive prediction. To enable comparison across sessions with different trial counts, we normalized flow by dividing by the number of trials and target-area dimensionality:

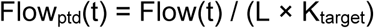

where L is the number of trials and K_target_ is the dimensionality of the target-area projection (K = 1 for scalar feature codes). To account for the positive bias expected when adding predictors to a regression model, we subtracted an approximate complexity baseline (Wilks):

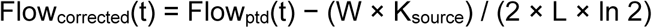

where W is the number of lag bins (i.e., the number of source-area predictors) and K_source_ is the source-area dimensionality. This correction removes the expected positive bias under the null hypothesis of no true predictive relationship. We computed corrected flow in both directions.

### Permutation testing and across-session inference for flow

To assess significance while controlling for condition structure and spurious cross-area correlations, we used permutation-based null distributions that preserve within-area temporal structure while breaking trial-to-trial coupling across areas. For each session, we generated a stratified trial-shuffle null with 500 permutations. Trial identities were permuted within task-condition strata, breaking the trial-to-trial correspondence between areas while preserving condition structure. For each permutation and time bin, we recomputed flow using the same lagged regression procedure. One-sided p-values were computed as p(t) = (1 + #{null(t) ≥ obs(t)}) / (1 + #{null valid(t)}).

We summarized flow time courses across sessions as mean ± SEM and computed net flow as the difference between the two directions. To assess significance of the across-session mean net-flow difference at each time bin, we constructed an empirical group null from per-session permutation samples: for each of 4,096 group replicates, we sampled one permutation draw per session, formed the per-session null net-flow difference, and averaged across sessions. To reduce sensitivity of group-level inference to bin-to-bin noise, we applied a uniform moving-average smoothing of width 20 ms to the observed mean net-flow time course and to each group-null replicate before computing group p-values. Group p-values were computed one-sided as the fraction of group-null means exceeding the observed group mean (with a +1 correction), and bins with p < 0.05 were marked in Fig. 5E,F and Fig. S7B,C,E,F.

## Supplemental Information

**Figure S1. Task outline and behavioral analyses for monkeys performing the rapid categorization task with horizontal targets. (A)** Trial structure of the rapid categorization task, such that one color target is positioned in the receptive field of recorded neurons (left visual field). Monkeys initiate each trial with central fixation and maintain fixation as motion stimuli and saccade targets appear in the periphery. After the fixation point disappears (Go Cue), the monkey must saccade to a color target within 500ms. However, the motion stimulus only becomes informative after a variable Gap Time (GT) following the Go Cue (see Methods). The positions of the color targets are randomized every trial. The category identity of the six motion directions are shown in the bottom-right inset, with center directions labeled. **(B)** Monkeys’ probability of fully committing to the red category target when presented with each direction stimulus, across all recording sessions, across all processing times. **(C)** Monkeys’ categorization accuracy quantified across a range of processing times (PT); accuracy quantified as the fraction of trials the monkey responded correctly. Shaded areas indicate SEM calculated from binomial statistics (see Methods). **(D)** Distribution of processing times of correct (gray bars) and incorrect trials (orange lines) from both monkeys.

**Figure S2. High percentages of single neurons in LIP, FEF, and SC are selective for category and direction of motion stimulus, configuration of targets, and direction of saccade.** (A) For each monkey and brain region, a bar graph representing the percent of single neurons significantly modulated by the following factors: stimulus category, stimulus direction, spatial target configuration, saccade direction, or a combination of features (also see Table S1). Significance was determined as Bonferroni-corrected p<0.05/8 by an ANOVA model fit for each task orientation at four time points relative to stimulus onset.

**Figure S3. Visual Receptive Fields and Saccadic Response Fields are concentrated in the left visual hemifield, contralateral to recording sites in FEF, LIP, and SC. (A)** Visual receptive fields of single neurons and multi-unit clusters from both monkeys, as determined by responses during Memory-Guided Saccade (MGS). Polar histogram includes the preferred MGS target locations, as determined by spike count by direction vector sums, of units selective by one-way ANOVA (p < 0.01) during the time window 0 to 300ms following target onset (Monkey M: FEF n = 291 units, LIP n = 747, SC n = 95; Monkey S: FEF n = 36, LIP n = 191, SC n = 70). **(B)** same as **a,** but for time window-300 to 0ms preceding MGS saccade onset (Monkey M: FEF n = 486, LIP n = 745, SC n = 115; Monkey S: FEF n = 142, LIP n = 179, SC n = 94).

**Figure S4. Direction and category selectivity are distributed at different time scales across LIP, FEF, and SC in the horizontal-target task orientation. (A)** Average single neuron ROC-based category tuning index (rCTI) by brain region (far left panel) and individual single neuron rCTIs arranged in rows as a heatmap (middle left panel) for Monkey M. The same format is used on the right panels to display results of single neurons for Monkey S. Only stimulus (category and direction) selective neurons as determined by ANOVA are shown. Shaded areas indicate SEM across single neurons. **(B)** Accuracy of linear support vector machines (SVMs) trained to predict the category of stimuli using activity from pseudopopulations constructed for each area. Left panel denotes the time course of SVM accuracy for Monkey M. Middle panel denotes the time course for Monkey S. Correct horizontal-target orientation trials in which the animal had at least 200ms of processing time, and the motion stimulus was placed above fixation were used for these analyses. Shaded intervals represent ± one standard deviation across pseudopopulations. Right panel shows scatter plots of the latency of SVMs for each pseudopopulation, defined as the first time point the SVM crossed an accuracy threshold of halfway between chance and the maximum accuracy attained by the SVM, for five consecutive time points (25ms). Box plots denote mean ± one standard deviation. Gray shading denotes neurons from Monkey M. **(C)** The same layout as **b,** but displaying results for SVMs trained to predict the direction of motion stimuli using activity of pseudopopulations for each area. Correct trials in which the animal had at least 200ms of processing time, and the motion stimulus was placed above the point of fixation and saccade targets were horizontally positioned were used for these analyses. Scatter plots depict the latency. *** denotes p<0.001, ** denotes p<0.01, subsampled permutation test.

**Figure S5. SVM decoding latency calculations with increasing pseudopopulation size. (A)** Mean category decoding latency of each monkey and brain area, calculated after 200 decoding runs (100 per direction-independent category decoding set; see Methods) at each pseudopopulation size, as decoder pseudopopulation size increases from 50 to the minimum available number of single neurons and multi-unit clusters, across the three areas. Latency is defined as the first time point the SVM crossed an accuracy threshold of halfway between chance and the maximum accuracy attained by the SVM, for five consecutive time points (25ms). **(B-D)**, same as **a**, but after 100 decoding runs of motion direction within green category, target configuration, and saccade direction, respectively.

**Figure S6. SC leads the encoding of target configuration, while core oculomotor output areas lead the encoding of saccade direction in the vertical-target task orientation. (A)** Accuracy of linear support vector machines (SVMs) trained to predict the target configuration (e.g. target color in RF) using activity from pseudopopulations constructed for each area. Left panel denotes the time course of SVM accuracy over time for Monkey M. Middle panel denotes the time course for Monkey S. Correct trials in which the stimulus was placed to the left of fixation, and targets were positioned vertically, were used. Shaded intervals represent ± one standard deviation across pseudopopulations. Right panel shows scatter plots of the latency of SVMs for each pseudopopulation, defined as the first time point the SVM crossed an accuracy threshold of halfway between chance and the maximum accuracy attained by the SVM, for five consecutive time points (25ms). Box plots denote mean ± one standard deviation. Gray shading denotes neurons from Monkey M. **(B)** The same layout as **a**, but displaying results for SVMs trained to predict the monkey’s saccade direction from both correct and incorrect trials, using activity of pseudopopulations for each area. *** denotes p<0.001, ** denotes p<0.01, * denotes p<0.05, subsampled permutation test.

**Figure S7. Single-trial onset latency and directed flow analyses for SC–LIP and FEF–SC area pairs. (A,B,C)** Monkey M. **(D,E,F)** Monkey S. **(A,D)** Single-trial onset latencies comparing SC versus LIP for category (left, stimulus-aligned) and saccade direction (right, saccade-aligned). Format follows Fig. 5C,D. Onset is defined as the first time signed evidence exceeded a baseline-derived threshold (μ + 4σ) for 5 consecutive bins; only trials with defined onsets in both areas were included. Dashed line: y = x. Large markers indicate the mean latency for each pair. Mean latency differences (LIP − SC) were: Monkey M category: −7.5 ms (N = 246, p = 0.412), saccade: −2.7 ms (N = 332, p = 0.669); Monkey S category: −17.1ms (N = 176, p = 0.137), saccade: 40.0 ms (N = 600, p < 0.001). P-values are two-sided sign-flip permutation tests on paired latency differences. SC and LIP showed no reliable onset latency difference for category in either monkey or for saccade direction in Monkey M; in Monkey S, LIP saccade onset significantly lagged SC. **(B,E)** Directed predictability gain (bits) between LIP and SC. Top: category (stimulus-aligned, 80 ms lag window). Bottom: saccade direction (saccade-aligned, 30 ms lag window). Format follows Fig. 5E,F. Curves show mean ± SEM across sessions; line color indicates the source area (red: LIP→SC, purple: SC→LIP). Colored dots mark time bins where one direction significantly exceeded the reverse under a stratified trial-shuffle null (p < 0.05); dot color indicates which direction was larger. **(C,F)** Directed predictability gain between FEF and SC for saccade direction (saccade-aligned, 30 ms lag window). Line color indicates the source area (blue: FEF→SC, purple: SC→FEF).

**Figure S8. Infusion of phosphate-buffered saline (PBS) into FEF does not significantly impair or delay categorization performance. (A),** Categorization performance in the RC task with vertically positioned targets (e.g. motion stimulus positioned within inactivated field, IF), is not impaired after saline infusion. Left panel displays the animal’s behavioral performance and biases quantified as a function of processing time for control and PBS infusion sessions, following the same format as **Fig 6C**. Dashed orange lines reflect the animal’s accuracy during inactivation. Shaded areas represent 95% confidence intervals calculated using binomial statistics. Right panel shows bar plots of the animal’s latency, defined as the time point the animal’s accuracy crossed 0.75 for 150 consecutive time bins. Error bars denote ±1SD across bootstrap iterations. n.s. denotes p>0.05 from the bootstrap percentile test. ‘C’ denotes control behavior, and ‘PBS’ denotes behavior after saline infusion. **(B),** The same format as **a,** quantifying performance in the RC task with horizontal targets and a vertically positioned motion stimulus.

**Table S1: Selectivities of single neurons and multi-unit clusters across FEF, LIP, and SC. (A),** Total counts of single neurons in each monkey and brain area (left), percentages of neurons selective for each task feature and saccade (category of visual motion stimulus, direction of visual motion stimulus, direction of saccade, and configuration of targets; see Figure S2). The right column depicts the percent of single neurons selective for at least one task feature or saccade direction. Significant feature selectivity was determined as Bonferroni-corrected p<0.05/8 by an ANOVA model fit for each task orientation at four time points relative to stimulus onset. **(B),** Same as **a,** but for multi-unit clusters.

**Table S2: Stimulus motion category and direction encoding latency statistics during vertical-target orientation trials across LIP, FEF, and SC. (A),** Stimulus-aligned latencies for stimulus category pseudopopulation SVM decoders (Fig. 3B; vertical-target orientation). Latencies were determined as the time at which SVM crossed an accuracy threshold of halfway between chance and the maximum accuracy attained by the SVM, for five consecutive time points (25ms). Significance of cross-area latency differences (right panel) as determined by subsampled permutation tests (20,000 permutations). **(B),** Same as **a,** but for stimulus-aligned latencies of stimulus motion direction pseudopopulation SVM decoders (Fig. 3C).

**Table S3: Stimulus motion category and direction encoding latency statistics during horizontal-target orientation trials across LIP, FEF, and SC. (A),** Stimulus-aligned latencies for stimulus category pseudopopulation SVM decoders (Fig. S4A; vertical-target orientation). Latencies were determined as the time at which SVM crossed an accuracy threshold of halfway between chance and the maximum accuracy attained by the SVM, for five consecutive time points (25ms). Significance of cross-area latency differences (right panel) as determined by subsampled permutation tests (20,000 permutations). **(B),** Same as **a,** but for stimulus-aligned latencies of stimulus motion direction pseudopopulation SVM decoders (Fig. S4B).

**Table S4: Saccade target configurations and saccade direction encoding latency statistics during horizontal-target orientation trials across LIP, FEF, and SC. (A),** Target onset-aligned latencies for pseudopopulation-based SVMs decoding (left/right) saccade target configuration (Fig. 4A; horizontal-target orientation). Latencies were determined as the time at which SVM crossed an accuracy threshold of halfway between chance and the maximum accuracy attained by the SVM, for five consecutive time points (25ms). Significance of cross-area latency differences (right panel) as determined by subsampled permutation tests (20,000 permutations). **(B),** Same as **a,** but for saccade onset-aligned latencies of SVMs decoding direction of saccadic choice (Fig. 4B).

**Table S5: Saccade target configurations and saccade direction encoding latency statistics during vertical-target orientation trials across LIP, FEF, and SC. (A),** Target onset-aligned latencies for pseudopopulation-based SVMs decoding (up/down) saccade target configuration (Fig. S6A). Latencies were determined as the time at which SVM crossed an accuracy threshold of halfway between chance and the maximum accuracy attained by the SVM, for five consecutive time points (25ms). Significance of cross-area latency differences (right panel) as determined by subsampled permutation tests (20,000 permutations). **(B),** Same as **a,** but for saccade onset-aligned latencies of SVMs decoding direction of saccadic choice (Fig. S6B).

**Table S6: Details of inactivation experiments in two monkeys.** Each row refers to an individual session, and each column provides details for that session. The columns describe: “Monkey” for the monkey in which the experiment was performed, “Exp.” the experiment number for that animal, “Treatment” for whether muscimol or saline was infused in that experiment, “Concentr.” for the concentration of muscimol or saline, “Inj. Vol.” for the total injection volume, “# RCT Trials (vertical orientation)” for the total number of attempted trials of the RC task with saccade targets arranged vertically and the stimulus positioned in the inactivated field, “# RCT Trials (horizontal orientation)” for the total number of attempted trials for the RC task with saccade targets arranged horizontally (with one target positioned inside the inactivated field) and the stimulus positioned above the fixation point.

