## Supplemental Information for "Frontal Eye Field Leads a Distributed Oculomotor Circuit for Abstract Categorical Decisions"

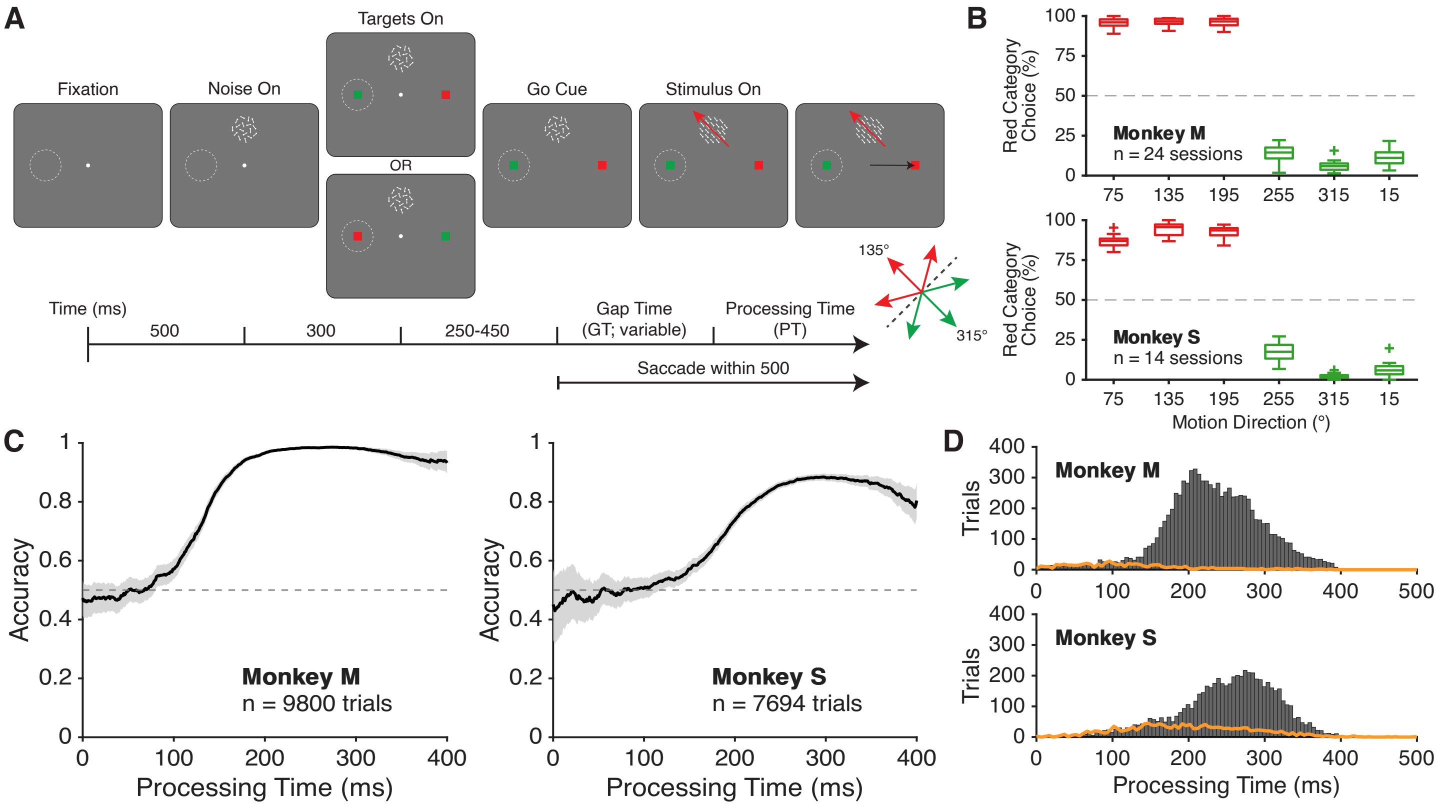


**Figure S1. Task outline and behavioral analyses for monkeys performing the rapid categorization task with horizontal targets. (A),** Trial structure of the rapid categorization task, such that one color target is positioned in the receptive field of recorded neurons (left visual field). Monkeys initiate each trial with central fixation and maintain fixation as motion stimuli and saccade targets appear in the periphery. After the fixation point disappears (Go Cue), the monkey must saccade to a color target within 500ms. However, the motion stimulus only becomes informative after a variable Gap Time (GT) following the Go Cue (see Methods). The positions of the color targets are randomized every trial. The category identity of the six motion directions are shown in the bottom-right inset, with center directions labeled. **(B)**, Monkeys' behavioral performance for each direction across all recording sessions, across all processing times. **(C)**, Monkeys' categorization accuracy quantified across a range of processing times (PT); accuracy quantified as fraction of trials the monkey responded correctly. Shaded areas indicate SEM calculated from binomial statistics (see Methods). **(D)**, Distribution of processing times of correct (gray bars) and incorrect trials (orange lines) from both monkeys.

**A**

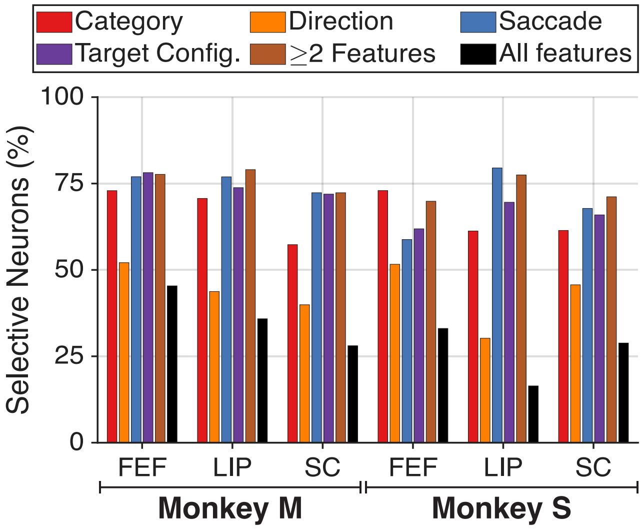


**Figure S2. High percentages of single neurons in LIP, FEF, and SC are selective for category and direction of motion stimulus, configuration of targets, and direction of saccade. (A),** For each monkey and brain region, a bar graph representing the percent of single neurons significantly modulated by the following factors: stimulus category, stimulus direction, spatial target configuration, saccade direction, or a combination of features (also see Table S1). Significance was determined as Bonferroni-corrected $p<0.05/8$ by an ANOVA model fit for each task orientation at four time points relative to stimulus onset.

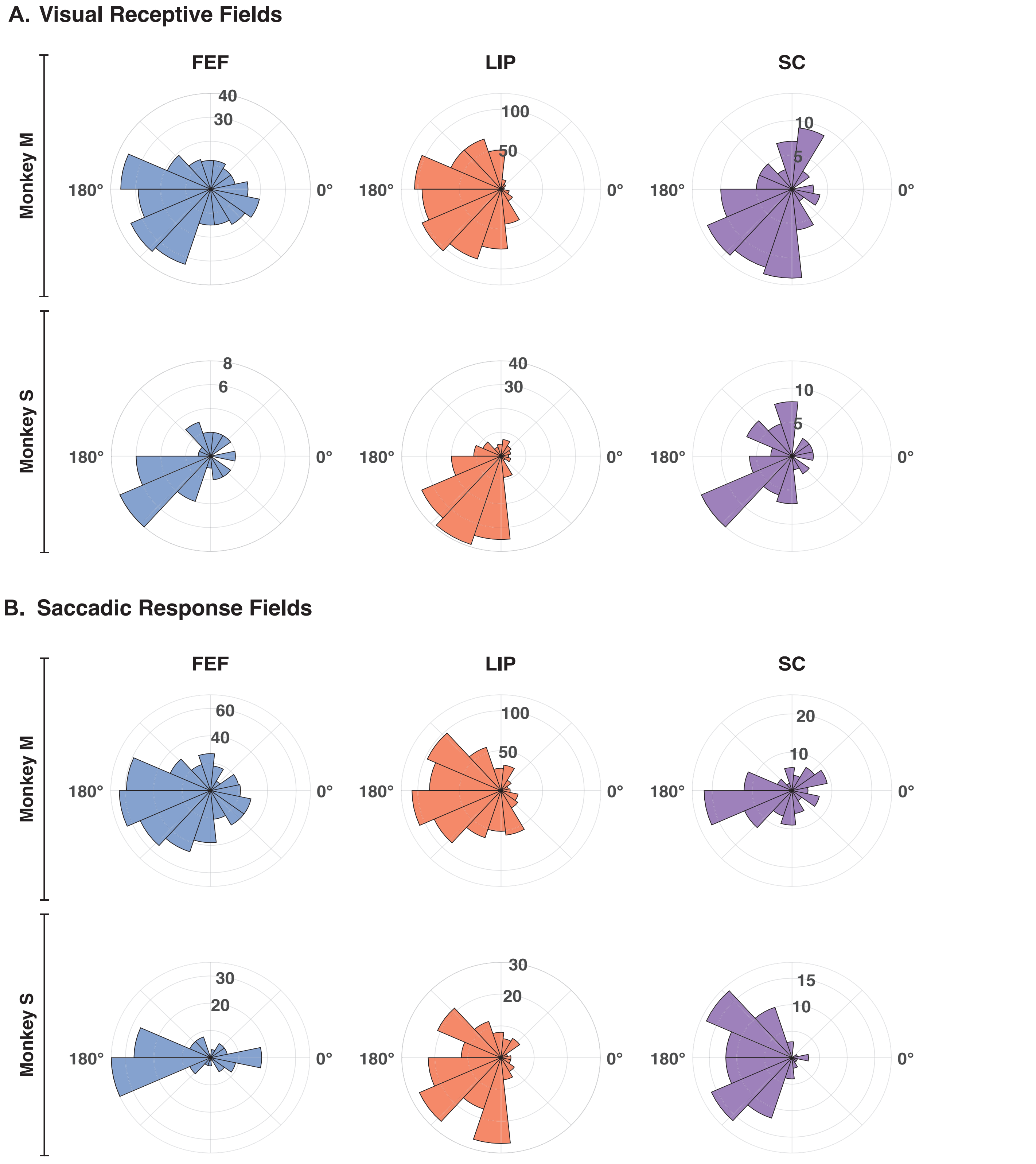


**Figure S3: Visual receptive fields and saccadic response fields are concentrated in the left visual hemifield, contralateral to recording sites in FEF, LIP, and SC. (A),** Visual receptive fields of single neurons and multi-unit clusters from both monkeys, as determined by responses during Memory-Guided Saccade (MGS). Polar histogram includes the preferred MGS target locations, as determined by spike count by direction vector sums, of units selective by one-way ANOVA ($p < 0.01$) during the time window 0 to 300ms following target onset (Monkey M: FEF n = 291 units, LIP n = 747, SC n = 95; Monkey S: FEF n = 36, LIP n = 191, SC n = 70). **(B),** same as **a,** but for time window -300 to 0ms preceding MGS saccade onset (Monkey M: FEF n = 486, LIP n = 745, SC n = 115; Monkey S: FEF n = 142, LIP n = 179, SC n = 94).

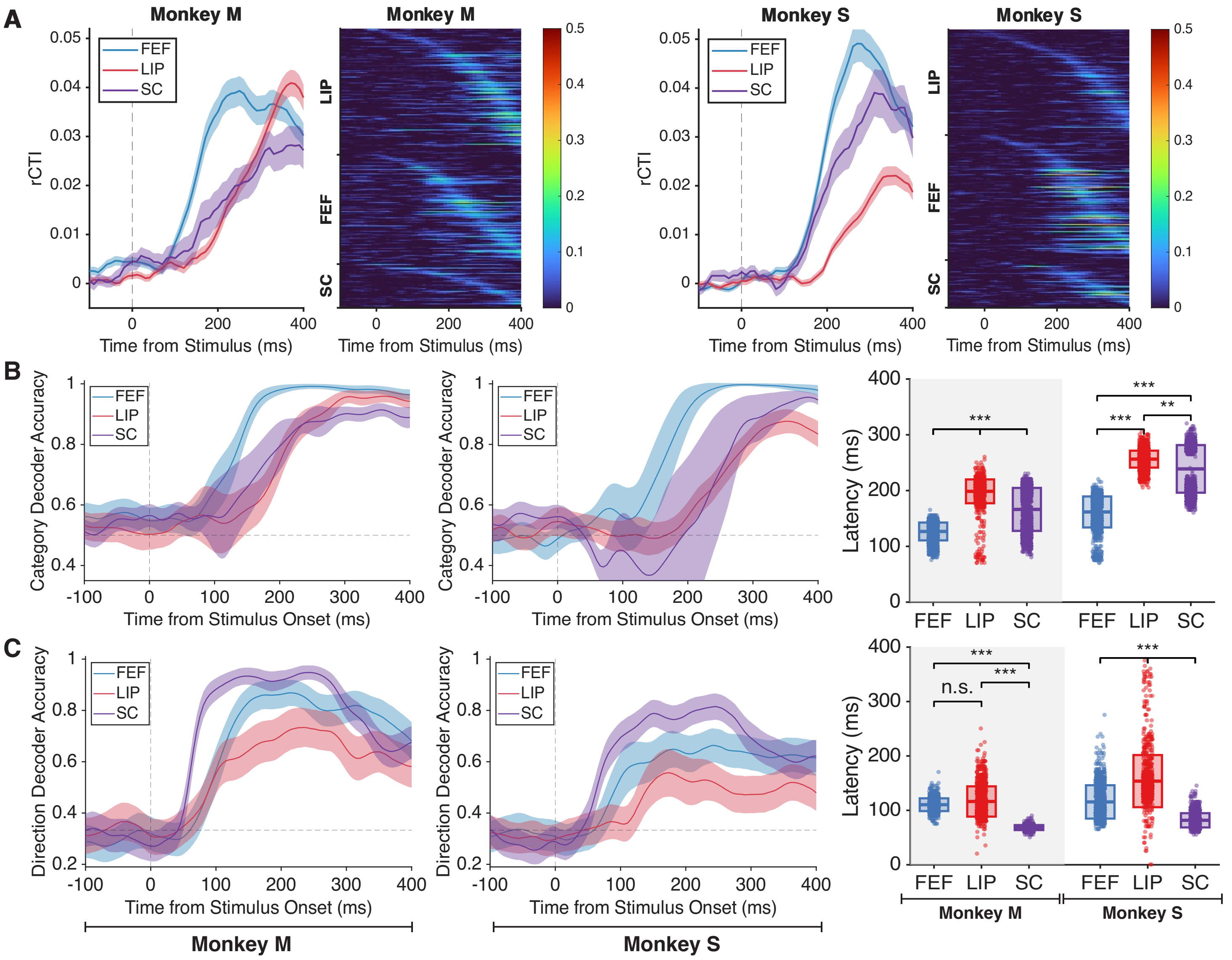


**Figure S4. Direction and category selectivity are distributed at different time scales across LIP, FEF, and SC in the horizontal-target task orientation. (A),** Average single neuron ROC-based category tuning index (rCTI) by brain region (far left panel) and individual single neuron rCTIs arranged in rows as a heatmap (middle left panel) for Monkey M. The same format is used on the right panels to display results of single neurons for Monkey S. Only stimulus (category and direction) selective neurons as determined by ANOVA are shown. Shaded areas indicate SEM across single neurons. **(B),** Accuracy of linear support vector machines (SVMs) trained to predict the category of stimuli using activity from pseudopopulations constructed for each area. Left panel denotes the time course of SVM accuracy for Monkey M. Middle panel denotes the time course for Monkey S. Correct horizontal-target orientation trials in which the animal had at least 200ms of processing time, and the motion stimulus was placed above fixation were used for these analyses. Shaded intervals represent ± one standard deviation across pseudopopulations. Right panel shows scatter plots of the latency of SVMs for each pseudopopulation, defined as the first time point the SVM crossed an accuracy threshold of halfway between chance and the maximum accuracy attained by the SVM, for five consecutive time points (25ms). Box plots denote mean ± one standard deviation. Gray shading denotes neurons from Monkey M. **(C),** The same layout as **b,** but displaying results for SVMs trained to predict the direction of motion stimuli using activity of pseudopopulations for each area. *** denotes p<0.001, ** denotes p<0.01, subsampled permutation test.

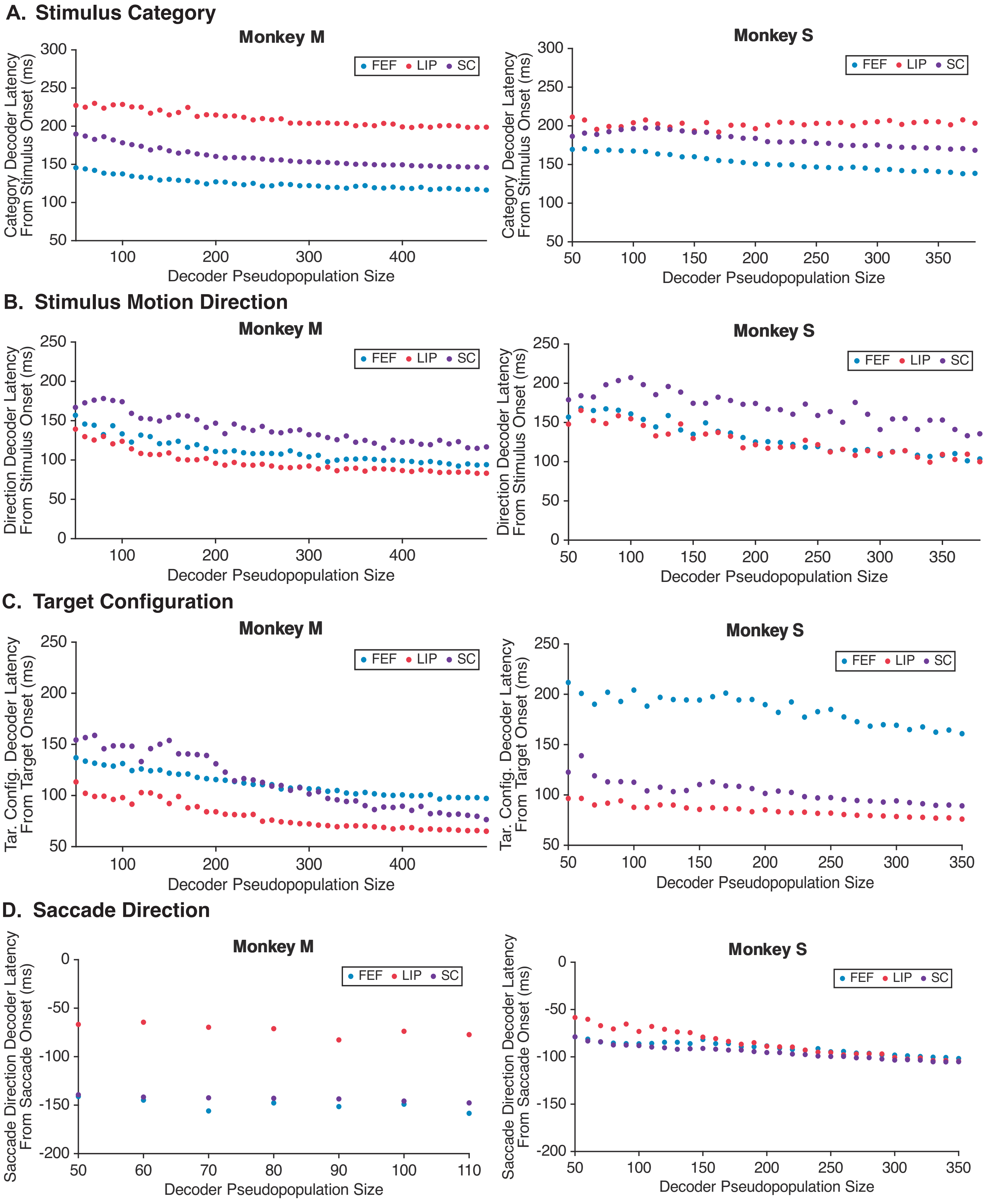


**Figure S5. SVM decoding latency calculations with increasing pseudopopulation size. (A),** Mean category decoding latency of each monkey and brain area, calculated after 200 decoding runs (100 per direction-independent category decoding set; see Methods) at each pseudopopulation size, as decoder pseudopopulation size increases from 50 to the minimum available number of single neurons and multi-unit clusters, across the three areas. Latency is defined as the first time point the SVM crossed an accuracy threshold of halfway between chance and the maximum accuracy attained by the SVM, for five consecutive time points (25ms). **(B-D),** same as **a,** but after 100 decoding runs of motion direction within green category, target configuration, and saccade direction, respectively.

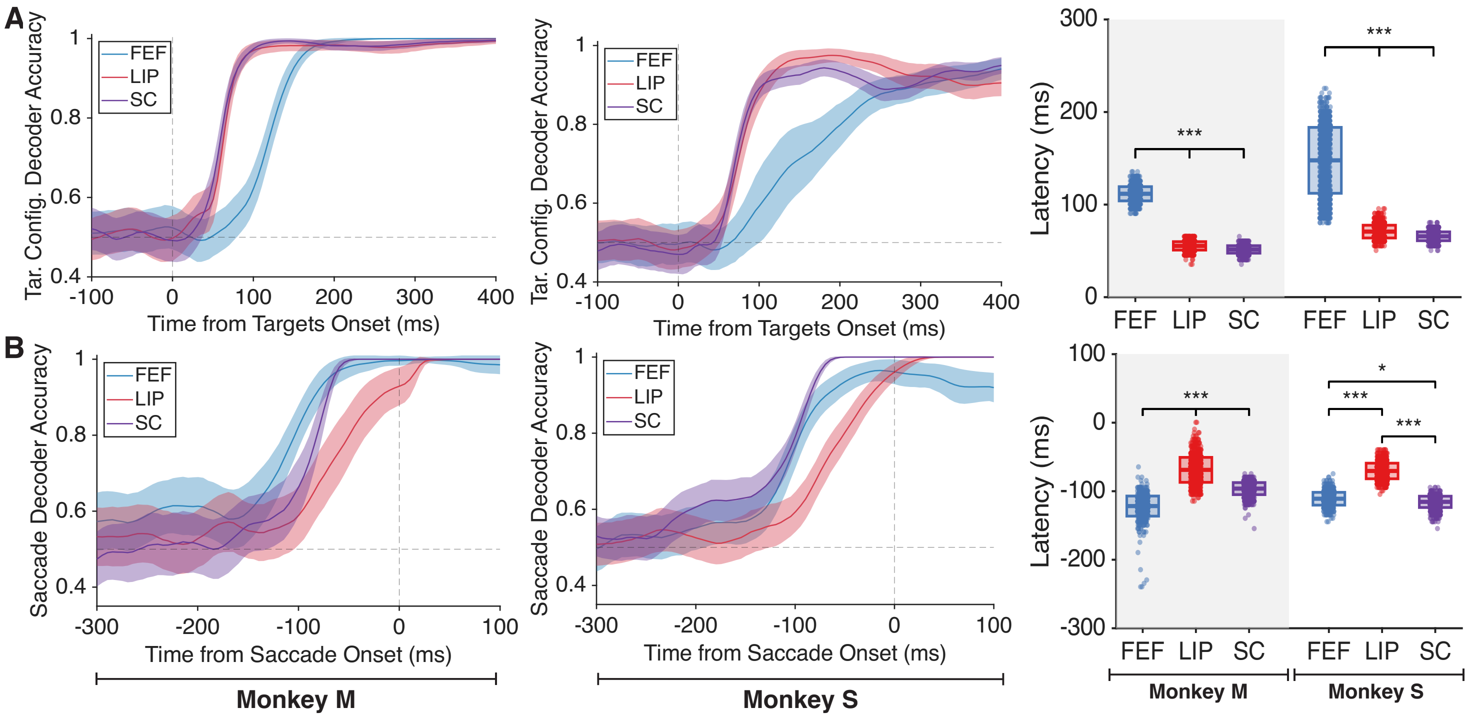


**Figure S6. SC leads the encoding of target configuration, while core oculomotor output areas lead the encoding of saccade direction in the vertical-target task orientation. (A),** Accuracy of linear support vector machines (SVMs) trained to predict the target configuration (e.g. target color in RF) using activity from pseudopopulations constructed for each area. Left panel denotes the time course of SVM accuracy over time for Monkey M. Middle panel denotes the time course for Monkey S. Correct trials in which the stimulus was placed to the left of fixation, and targets were positioned vertically, were used. Shaded intervals represent ± one standard deviation across pseudopopulations. Right panel shows scatter plots of the latency of SVMs for each pseudopopulation, defined as the first time point the SVM crossed an accuracy threshold of halfway between chance and the maximum accuracy attained by the SVM, for five consecutive time points (25ms). Box plots denote mean ± one standard deviation. Gray shading denotes neurons from Monkey M. **(B),** The same layout as **a**, but displaying results for SVMs trained to predict the monkey's saccade direction from both correct and incorrect trials, using activity of pseudopopulations for each area. *** denotes $p<0.001$, ** denotes $p<0.01$, * denotes $p<0.05$, subsampled permutation test.

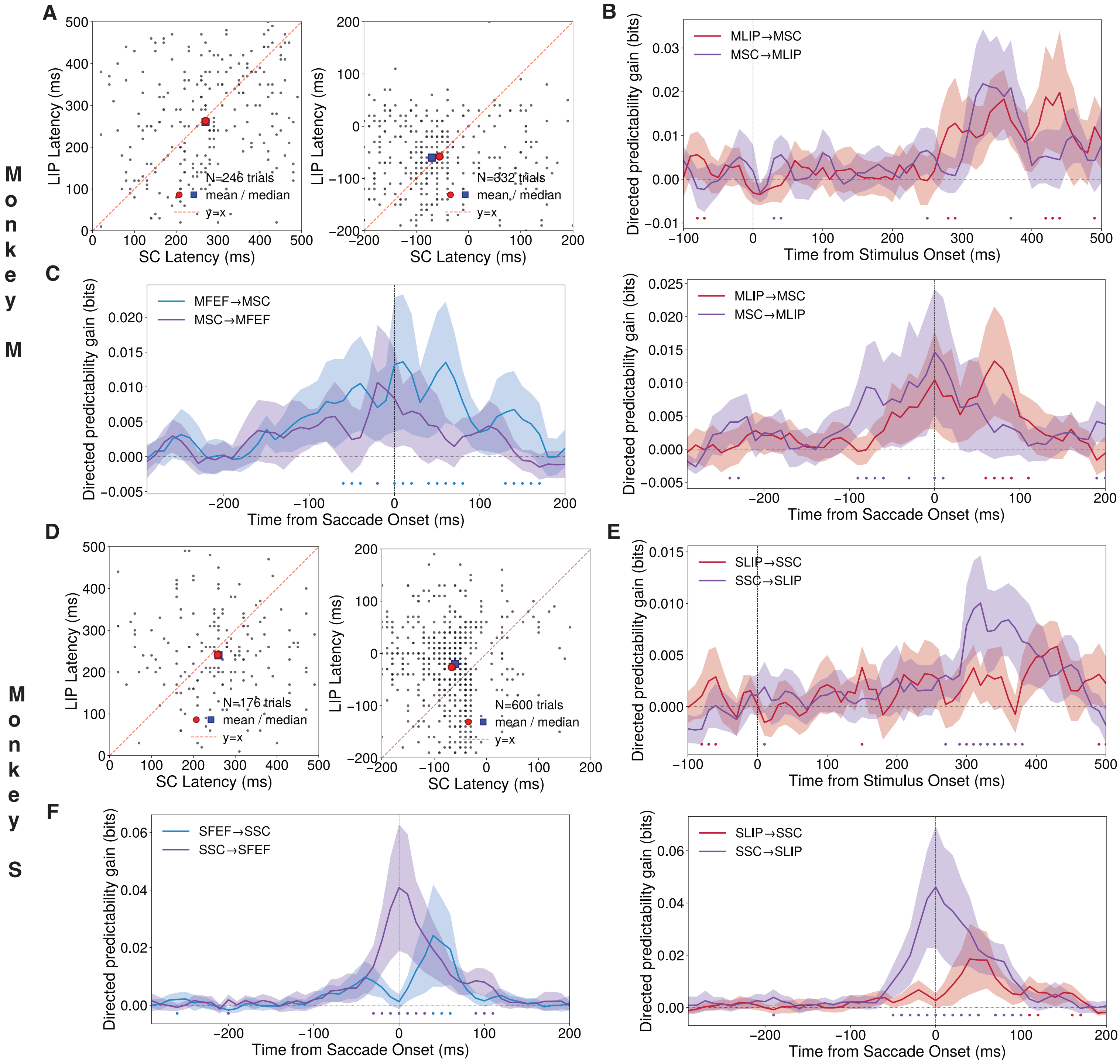


**Figure S7. Single-trial onset latency and directed flow analyses for SC–LIP and FEF–SC area pairs. (A,B,C)** Monkey M. **(D,E,F)** Monkey S. **(A,D)** Single-trial onset latencies comparing SC versus LIP for category (left, stimulus-aligned) and saccade direction (right, saccade-aligned). Format follows Fig. 5C,D. Onset is defined as the first time signed evidence exceeded a baseline-derived threshold ($\mu + 4\sigma$) for 5 consecutive bins; only trials with defined onsets in both areas were included. Dashed line: y = x. Large markers indicate the mean latency for each pair. Mean latency differences (LIP − SC) were: Monkey M category: −7.5 ms ($N = 246$, $p = 0.412$), saccade: −2.7 ms ($N = 332$, $p = 0.669$); Monkey S category: −17.1ms ($N = 176$, $p = 0.137$), saccade: 40.0 ms ($N = 600$, $p < 0.001$). P-values are two-sided sign-flip permutation tests on paired latency differences. SC and LIP showed no reliable onset latency difference for category in either monkey or for saccade direction in Monkey M; in Monkey S, LIP saccade onset significantly lagged SC. **(B,E)** Directed predictability gain (bits) between LIP and SC. Top: category (stimulus-aligned, 80 ms lag window). Bottom: saccade direction (saccade-aligned, 30 ms lag window). Format follows **Fig. 5E,F.** Curves show mean ± SEM across sessions; line color indicates the source area (red: LIP→SC, purple: SC→LIP). Colored dots mark time bins where one direction significantly exceeded the reverse under a stratified trial-shuffle null ($p < 0.05$); dot color indicates which direction was larger. **(C,F)** Directed predictability gain between FEF and SC for saccade direction (saccade-aligned, 30 ms lag window). Line color indicates the source area (blue: FEF→SC, purple: SC→FEF).

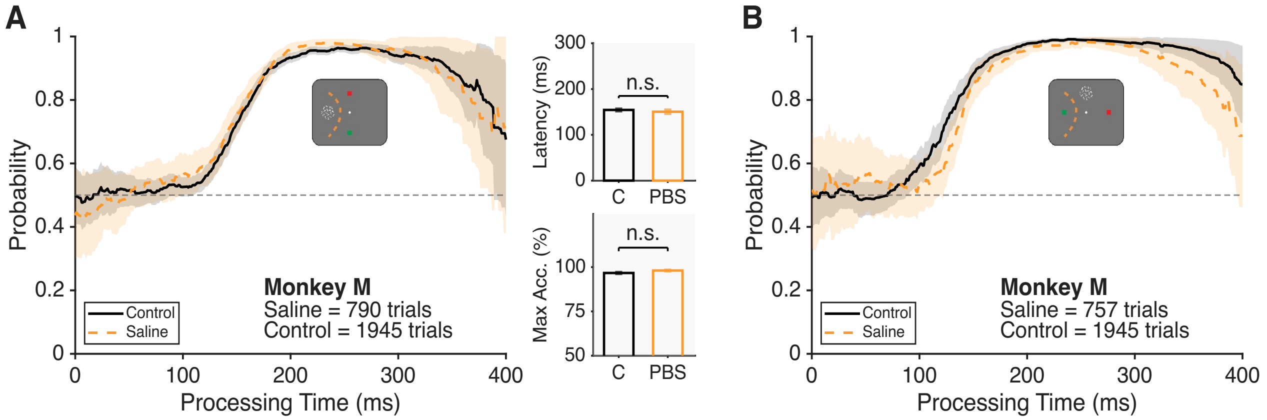


**Figure S8: Infusion of phosphate-buffered saline (PBS) into FEF does not significantly impair or delay categorization performance. (A),** Categorization performance in the RC task with vertically positioned targets (e.g. motion stimulus positioned within inactivated field, IF), is not impaired after saline infusion. Left panel displays the animal's behavioral performance and biases quantified as a function of processing time for control and PBS infusion sessions, following the same format as **Fig 6C**. Dashed orange lines reflect the animal's accuracy during inactivation. Shaded areas represent 95% confidence intervals calculated using binomial statistics. Right panel shows bar plots of the animal's latency, defined as the time point the animal's accuracy crossed 0.75 for 150 consecutive time bins. Error bars denote ±1SD across bootstrap iterations. n.s. denotes $p>0.05$ from the bootstrap percentile test. 'C' denotes control behavior, and 'PBS' denotes behavior after saline infusion. **(B),** The same format as **a**, quantifying performance in the RC task with horizontal targets and a vertically positioned motion stimulus.

**A. Single Neuron Selectivities**

| | Area | Total Count | Category-Selective | Direction-Selective | Saccade-Selective | Target-Selective | All Task-Modulated |
|---|---|---|---|---|---|---|---|
| **Monkey M** | FEF | 595 | 72.94% | 52.10% | 76.97% | 78.15% | 88.07% |
| | LIP | 672 | 70.68% | 43.75% | 76.93% | 73.81% | 88.10% |
| | SC | 253 | 57.31% | 39.92% | 72.33% | 71.94% | 83.00% |
| **Monkey S** | FEF | 614 | 72.96% | 51.63% | 58.79% | 61.89% | 86.81% |
| | LIP | 493 | 61.26% | 30.22% | 79.51% | 69.57% | 93.91% |
| | SC | 267 | 61.42% | 45.69% | 67.79% | 65.92% | 86.89% |

**B. Multi-unit Cluster Selectivities**

| | Area | Total Count | Category-Selective | Direction-Selective | Saccade-Selective | Target-Selective | All Task-Modulated |
|---|---|---|---|---|---|---|---|
| **Monkey M** | FEF | 417 | 73.62% | 42.93% | 77.46% | 75.78% | 88.73% |
| | LIP | 631 | 71.00% | 43.90% | 77.65% | 76.54% | 87.64% |
| | SC | 240 | 48.75% | 26.67% | 61.25% | 56.67% | 78.75% |
| **Monkey S** | FEF | 292 | 57.19% | 35.96% | 38.01% | 44.86% | 70.89% |
| | LIP | 214 | 54.21% | 23.36% | 73.36% | 57.01% | 87.85% |
| | SC | 118 | 42.37% | 44.92% | 54.24% | 49.15% | 77.97% |

**SI Appendix Table S1: Selectivities of single neurons and multi-unit clusters across FEF, LIP, and SC. (A),** Total counts of single neurons in each monkey and brain area (left), percentages of neurons selective for each task feature and saccade (category of visual motion stimulus, direction of visual motion stimulus, direction of saccade, and configuration of targets; see Figure S2). The right column depicts the percent of single neurons selective for at least one task feature or saccade direction. Significant feature selectivity was determined as Bonferroni-corrected $p<0.05/8$ by an ANOVA model fit for each task orientation at four time points relative to stimulus onset. **(B),** Same as **a,** but for multi-unit clusters.

**A. Stimulus Category Decoding: Vertical-Target Orientation (Stimulus-in)**

| | Area | Latency Mean | Latency STD | Runs Crossing Threshold | Latency Differences & Significance |
|---|---|---|---|---|---|
| **Monkey M** | FEF | 127.0 ms | 12.1 ms | n = 2000 | FEF leads LIP by 80.1 ms ($p < 0.0001$)<br>SC leads LIP by 51.2 ms, ($p < 0.0001$)<br>FEF leads SC by 28.9 ms, ($p < 0.0001$) |
| | LIP | 207.1 ms | 16.0 ms | n = 1999 | |
| | SC | 155.9 ms | 4.6 ms | n = 2000 | |
| **Monkey S** | FEF | 148.9 ms | 27.4 ms | n = 2000 | FEF leads LIP by 64.5 ms ($p < 0.0001$)<br>SC leads LIP by 34.3 ms, ($p < 0.0001$)<br>FEF leads SC by 30.2 ms, ($p < 0.0001$) |
| | LIP | 213.4 ms | 25.6 ms | n = 2000 | |
| | SC | 179.1 ms | 8.1 ms | n = 2000 | |

**B. Stimulus Motion Direction Decoding: Vertical-Target Orientation (Stimulus-in)**

| | Area | Latency Mean | Latency STD | Runs Crossing Threshold | Latency Differences & Significance |
|---|---|---|---|---|---|
| **Monkey M** | FEF | 104.1 ms | 16.7 ms | n = 1000 | LIP leads FEF by 10.6 ms ($p = 0.0002$)<br>LIP leads SC by 31.9 ms ($p < 0.0001$)<br>FEF leads SC by 21.3 ms ($p < 0.0001$) |
| | LIP | 93.5 ms | 9.2 ms | n = 1000 | |
| | SC | 125.4 ms | 20.6 ms | n = 1000 | |
| **Monkey S** | FEF | 116.0 ms | 26.6 ms | n = 1000 | LIP leads FEF by 7.7 ms ($p = 0.3762$)<br>LIP leads SC by 46.3 ms ($p = 0.0003$)<br>FEF leads SC by 38.6 ms ($p = 0.0004$) |
| | LIP | 108.3 ms | 54.5 ms | n = 999 | |
| | SC | 154.6 ms | 67.5 ms | n = 1000 | |

**SI Appendix Table S2: Stimulus motion category and direction encoding latency statistics during vertical-target orientation trials across LIP, FEF, and SC. (A),** Stimulus-aligned latencies for stimulus category pseudopopulation SVM decoders (Fig. 3B; vertical-target orientation). Latencies were determined as the time at which SVM crossed an accuracy threshold of halfway between chance and the maximum accuracy attained by the SVM, for five consecutive time points (25ms). Significance of cross-area latency differences (right panel) as determined by subsampled permutation tests (20,000 permutations). **(B),** Same as **a,** but for stimulus-aligned latencies of stimulus motion direction pseudopopulation SVM decoders (Fig. 3C).

**A. Stimulus Category Decoding: Horizontal-Target Orientation (Target-in)**

| | Area | Latency Mean | Latency STD | Runs Crossing Threshold | Latency Differences & Significance |
|---|---|---|---|---|---|
| **Monkey M** | FEF | 126.7 ms | 16.0 ms | n = 1992 | FEF leads LIP by 71.9 ms ($p < 0.0001$)<br>SC leads LIP by 32.4 ms ($p < 0.0001$)<br>FEF leads SC by 39.5 ms ($p < 0.0001$) |
| | LIP | 198.6 ms | 21.4 ms | n = 1977 | |
| | SC | 166.2 ms | 38.6 ms | n = 2000 | |
| **Monkey S** | FEF | 161.7 ms | 27.8 ms | n = 2000 | FEF leads LIP by 94.7 ms ($p < 0.0001$)<br>SC leads LIP by 17.5 ms ($p = 0.0085$)<br>FEF leads SC by 77.2 ms ($p < 0.0001$) |
| | LIP | 256.4 ms | 15.2 ms | n = 2000 | |
| | SC | 238.9 ms | 42.7 ms | n = 2000 | |

**B. Stimulus Motion Direction Decoding: Horizontal-Target Orientation (Target-in)**

| | Area | Latency Mean | Latency STD | Runs Crossing Threshold | Latency Differences & Significance |
|---|---|---|---|---|---|
| **Monkey M** | FEF | 110.4 ms | 11.7 ms | n = 998 | FEF leads LIP by 5.9 ms ($p = 0.1641$)<br>SC leads LIP by 47.9 ms ($p < 0.0001$)<br>SC leads FEF by 42.0 ms ($p < 0.0001$) |
| | LIP | 116.3 ms | 27.8 ms | n = 998 | |
| | SC | 68.4 ms | 4.6 ms | n = 1000 | |
| **Monkey S** | FEF | 115.5 ms | 30.7 ms | n = 1000 | FEF leads LIP by 38.2 ms ($p < 0.0001$)<br>SC leads LIP by 72.0 ms ($p < 0.0001$)<br>SC leads FEF by 33.8 ms ($p < 0.0001$) |
| | LIP | 153.7 ms | 47.9 ms | n = 999 | |
| | SC | 81.7 ms | 13.1 ms | n = 1000 | |

**SI Appendix Table S3: Stimulus motion category and direction encoding latency statistics during horizontal-target orientation trials across LIP, FEF, and SC. (A),** Stimulus-aligned latencies for stimulus category pseudopopulation SVM decoders (Fig. S4A; vertical-target orientation). Latencies were determined as the time at which SVM crossed an accuracy threshold of halfway between chance and the maximum accuracy attained by the SVM, for five consecutive time points (25ms). Significance of cross-area latency differences (right panel) as determined by subsampled permutation tests (20,000 permutations). **(B),** Same as **a,** but for stimulus-aligned latencies of stimulus motion direction pseudopopulation SVM decoders (Fig. S4B).

**A. Target Configuration Decoding: Horizontal-Target Orientation (Target-in)**

| | Area | Latency Mean | Latency STD | Runs Crossing Threshold | Latency Differences & Significance |
|---|---|---|---|---|---|
| **Monkey M** | FEF | 96.8 ms | 7.5 ms | n = 1000 | LIP leads FEF by 31.9ms ($p < 0.0001$)<br>LIP leads SC by 12.3ms ($p < 0.0001$)<br>SC leads FEF by 19.6ms ($p < 0.0001$) |
| | LIP | 64.9 ms | 5.6 ms | n = 1000 | |
| | SC | 77.2 ms | 8.0 ms | n = 1000 | |
| **Monkey S** | FEF | 160.0 ms | 18.9 ms | n = 1000 | LIP leads FEF by 83.4ms ($p < 0.0001$)<br>LIP leads SC by 12.0ms ($p < 0.0001$)<br>SC leads FEF by 71.4ms ($p < 0.0001$) |
| | LIP | 76.6 ms | 6.3 ms | n = 1000 | |
| | SC | 88.6 ms | 7.9 ms | n = 1000 | |

**B. Saccadic Response Direction Decoding: Horizontal-Target Orientation (Target-in)**

| | Area | Latency Mean | Latency STD | Runs Crossing Threshold | Latency Differences & Significance |
|---|---|---|---|---|---|
| **Monkey M** | FEF | -158.8ms | 38.9 ms | n = 1000 | FEF leads LIP by 81.0ms ($p < 0.0001$)<br>SC leads LIP by 68.4ms ($p < 0.0001$)<br>FEF leads SC by 12.6ms ($p = 0.0439$) |
| | LIP | -77.8ms | 35.5 ms | n = 1000 | |
| | SC | -146.2ms | 19.7 ms | n = 1000 | |
| **Monkey S** | FEF | -102.3ms | 9.9 ms | n = 1000 | LIP leads FEF by 2.3ms ($p = 0.2256$)<br>SC leads LIP by 1.1ms ($p = 0.4822$)<br>FEF leads SC by 3.4ms ($p = 0.0428$) |
| | LIP | -104.6ms | 9.7 ms | n = 1000 | |
| | SC | -105.7ms | 6.5 ms | n = 1000 | |

**SI Appendix Table S4: Saccade target configurations and saccade direction encoding latency statistics during horizontal-target orientation trials across LIP, FEF, and SC. (A),** Target onset-aligned latencies for pseudopopulation-based SVMs decoding (left/right) saccade target configuration (Fig. 4A; horizontal-target orientation). Latencies were determined as the time at which SVM crossed an accuracy threshold of halfway between chance and the maximum accuracy attained by the SVM, for five consecutive time points (25ms). Significance of cross-area latency differences (right panel) as determined by subsampled permutation tests (20,000 permutations). **(B),** Same as **a,** but for saccade onset-aligned latencies of SVMs decoding direction of saccadic choice (Fig. 4B).

**A. Target Configuration Decoding: Vertical-Target Orientation (Stimulus-in)**

| | Area | Latency Mean | Latency STD | Runs Crossing Threshold | Latency Differences & Significance |
|---|---|---|---|---|---|
| **Monkey M** | FEF | 111.7 ms | 7.7 ms | n = 1000 | LIP leads FEF by 56.3ms (p < 0.0001) SC leads LIP by 3.8ms (p < 0.0001) SC leads FEF by 60.1ms (p < 0.0001) |
| | LIP | 55.4 ms | 4.5 ms | n = 1000 | |
| | SC | 51.6 ms | 4.0 ms | n = 1000 | |
| **Monkey S** | FEF | 147.8 ms | 35.6 ms | n = 1000 | LIP leads FEF by 77.0ms (p < 0.0001) SC leads LIP by 5.0ms (p = 0.00005) SC leads FEF by 82.0ms (p < 0.0001) |
| | LIP | 70.8 ms | 6.9 ms | n = 1000 | |
| | SC | 65.8 ms | 4.8 ms | n = 1000 | |

**B. Saccadic Response Direction Decoding: Vertical-Target Orientation (Stimulus-in)**

| | Area | Latency Mean | Latency STD | Runs Crossing Threshold | Latency Differences & Significance |
|---|---|---|---|---|---|
| **Monkey M** | FEF | -121.7ms | 14.7 ms | n = 1000 | FEF leads LIP by 52.9ms (p < 0.0001) SC leads LIP by 27.4ms (p < 0.0001) FEF leads SC by 25.5ms (p < 0.0001) |
| | LIP | -68.8ms | 18.2 ms | n = 1000 | |
| | SC | -96.2ms | 8.8 ms | n = 1000 | |
| **Monkey S** | FEF | -111.0ms | 9.3 ms | n = 1000 | FEF leads LIP by 40.5ms (p < 0.0001) SC leads LIP by 45.0ms (p < 0.0001) SC leads FEF by 4.5ms (p = 0.0109) |
| | LIP | -70.5ms | 11.4 ms | n = 1000 | |
| | SC | -115.5ms | 8.2 ms | n = 1000 | |

**SI Appendix Table S5: Saccade target configurations and saccade direction encoding latency statistics during vertical-target orientation trials across LIP, FEF, and SC. (A),** Target onset-aligned latencies for pseudopopulation-based SVMs decoding (up/down) saccade target configuration (Fig. S6A). Latencies were determined as the time at which SVM crossed an accuracy threshold of halfway between chance and the maximum accuracy attained by the SVM, for five consecutive time points (25ms). Significance of cross-area latency differences (right panel) as determined by subsampled permutation tests (20,000 permutations). **(B),** Same as **a,** but for saccade onset-aligned latencies of SVMs decoding direction of saccadic choice (Fig. S6B).

| | Exp. | Treatment | Concentr. (μg/μL) | Inj. Vol. (μL) | #RCT trials (vertical orientation) | #RCT trials (horizontal orientation) |
|---|---|---|---|---|---|---|
| **Monkey M** | 1 | Muscimol | 5 | 1.50 | 775 | 490 |
| | 2 | Muscimol | 5 | 2.00 | 284 | 254 |
| | 3 | Muscimol | 5 | 2.00 | 410 | 442 |
| | 4 | Muscimol | 5 | 2.00 | 424 | 461 |
| | 5 | Muscimol | 5 | 2.33 | 443 | 387 |
| | 6 | Muscimol | 5 | 2.33 | 480 | 386 |
| | 7 | Muscimol | 5 | 2.33 | 459 | 462 |
| | 8 | Muscimol | 5 | 2.33 | 497 | 410 |
| | 9 | Saline | 9 | 2.33 | 396 | 373 |
| | 10 | Saline | 9 | 2.33 | 394 | 384 |
| **Monkey S** | 1 | Muscimol | 5 | 2.17 | 647 | 414 |
| | 2 | Muscimol | 5 | 2.17 | 671 | 481 |
| | 3 | Muscimol | 5 | 2.08 | 316 | 0 |
| | 4 | Muscimol | 5 | 2.08 | 352 | 0 |
| | 5 | Muscimol | 5 | 2.75 | 471 | 0 |

**SI Appendix Table S6: Details of inactivation experiments in two monkeys.** Each row refers to an individual session, and each column provides details for that session. The columns describe: "Monkey" for the monkey in which the experiment was performed, "Exp." the experiment number for that animal, "Treatment" for whether muscimol or saline was infused in that experiment, "Concentr." for the concentration of muscimol or saline, "Inj. Vol." for the total injection volume, "# RCT Trials (vertical orientation)" for the total number of attempted trials of the RC task with saccade targets arranged vertically and the stimulus positioned in the inactivated field, "# RCT Trials (horizontal orientation)" for the total number of attempted trials for the RC task with saccade targets arranged horizontally (with one target positioned inside the inactivated field) and the stimulus positioned above the fixation point.
